# Predicting immunotherapy response in advanced bladder cancer: a meta-analysis of six independent cohorts

**DOI:** 10.1101/2024.04.18.589711

**Authors:** Lilian Marie Boll, Sergio Vázquez Montes de Oca, Marta E. Camarena, Robert Castelo, Joaquim Bellmunt, Júlia Perera-Bel, M. Mar Albà

**Author notes:** co-corresponding authors (JB) (JP-B) (MMA). co-first authors (Lilian Marie Boll, Sergio Vázquez Montes de Oca).

## Abstract

Advanced bladder cancer patients show very variable responses to immune checkpoint inhibitors (ICIs) and effective strategies to predict response are still lacking. Here we integrate mutation and gene expression data from 707 advanced bladder cancer patients treated with anti-PD-1/anti-PD-L1 to build highly accurate predictive models. We find that, in addition to tumor mutational burden (TMB), enrichment in the APOBEC mutational signature, and the abundance of pro-inflammatory macrophages, are major factors associated with the response. Paradoxically, patients with high immune infiltration do not show an overall better response. We show that this can be explained by the activation of immune suppressive mechanisms in a large portion of these patients. In the case of non-immune-infiltrated cancer subtypes, we uncover specific variables likely to be involved in the response. Our findings provide novel information for advancing precision medicine in patients with advanced bladder cancer treated with immunotherapy.

## INTRODUCTION

Immune checkpoint inhibitors (ICI) have been a major breakthrough in the treatment of advanced bladder cancer ^1,2^. ICI can result in partial or even complete remission of the tumor, but it is only effective in a subset of the patients. Understanding which are the factors that underlie the response to ICI is key to improve our ability to predict response and to develop better therapeutic instruments^3,4^. While the number of somatic mutations in the tumor, or tumor mutational burden (TMB), is positively associated with ICI response, TMB alone has a moderate predictive power ^5,6^. Another biomarker with conflicting results is the expression of the programmed cell death-ligand 1 (PD-L1) together with its receptor, programmed cell death protein (PD-1), targeted by ICI drugs ^1,2,7,8^. Immune players that have been associated with the response include the abundance of CD8+ T cells, positively associated with the response in inflamed tumors, and the transforming growth factor β (TGF-β) signaling in fibroblasts, with a negative effect on the response to ICI ^9^.

Bladder cancer (BLCA) shows a high degree of heterogeneity, and five muscle-invasive subtypes have been defined in the TCGA urothelial bladder carcinoma cohort ^10,11,12^. The basal-squamous subtype is characterized by high expression of basal and stem-like markers, as well as immune markers. The luminal papillary subtype corresponds to tumors with papillary morphology and enriched in FGFR3 mutations. The luminal infiltrated subtype shows low tumor purity and muscle-related signatures. The luminal subtype is associated with high levels of uroplakins UPK1A and UPK2. Finally, the neuronal class is associated with high levels of neuronal differentiation and development genes. Whether these different subtypes show a similar or different response rate to immunotherapy is a matter of debate. The luminal infiltrated and basal squamous subtypes have a high proportion of infiltrated immune cells and this has been hypothesized to result in more effective responses to ICI ^10,13^. However, the results of a recent clinical trial have suggested that neuronal is the only subtype associated with a higher response ^14^.

Increasing our ability to predict the response to ICI is key to providing better treatments to patients. This can be achieved by building computational predictive models capable of integrating disparate sources of omics data ^15–17^. For these models to be robust, it is fundamental to gather data from a large number of patients. In order to address this challenge, we have assembled data from 707 tumors from six different cohorts of advanced BLCA patients treated with anti-PD-1/PD-L1. The models we have developed show high predictive accuracy and at the same time provide novel biological insights into the determinants of ICI response in the different subtypes.

## RESULTS

### Building an advanced BLA meta-cohort

We extracted clinical and omics data for a total of 707 advanced BLCA patients treated with anti-PD-L1/PD-1 (Figure 1). The patients were diagnosed with metastatic or locally advanced BLCA. Paired tumor-germline whole-exome sequencing (WES) data and tumor RNA sequencing (RNA-Seq) data were available for IMvigor210 ^9^, HdM-BLCA-1^18^, and SNY-2017 ^5^. For MIAO-2018 ^3^, WES data was collected. For the cohorts UC-GENOME ^19^ and UNC-108 ^16^, tumor RNA-Seq data, together with targeted DNA sequencing for a subset of patients, was available. To allow for a meaningful comparison of ICI therapy responders and non-responders, we considered the RECIST status of complete response (CR) and partial response (PR) as responders (class R) and PD (progressive disease) as non-responders (class NR). Stable disease (SD) and NE (not evaluable) were not considered for further analyses. We obtained a total of 466 patients classified as R or NR (163 and 303, respectively). For the majority of these patients (n=348, 75%) both mutation and tumor gene expression data was available.

**Figure 1.**
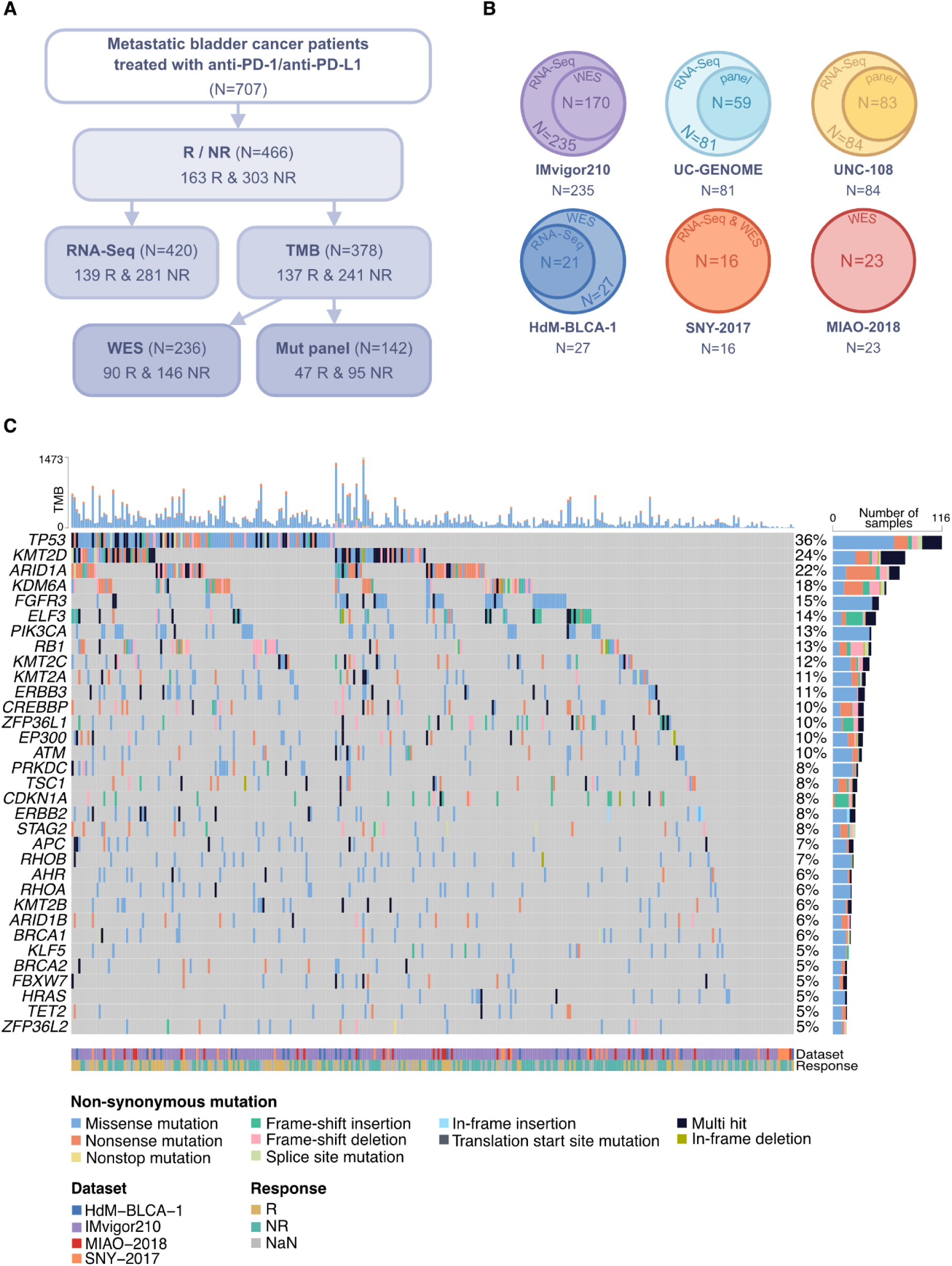
Overview of the analyzed sequencing data of six metastatic bladder cancer datasets. **A.** Number of patients with different types of data. Sequencing data of 707 patients was downloaded. Patients with a RECIST classification of partial or complete response to anti-PD-L1/PD-1 were considered responders (R) and progressive disease were considered non-responders (NR). Stable disease and non-evaluable were not included for further analysis. **B.** Of 466 patients with response to immunotherapy information, 348 patients have RNA-Seq data and a TMB estimate available. **C.** Most frequently mutated BLCA genes mutated in ≥5% of the samples over all four datasets with WES data (N=318). The X axis represents the patients from the different cohorts, the corresponding TMB is indicated. Mutations are classified in different types as indicated; multi hit describes cases where the gene carries more than one mutation in the patient. R: responders, NR: non-responders, Mut panel: DNA mutation panel.

Investigation of clinical and demographic variables indicated that having a low functional status (ECOG score ≥ 1) and presence of liver metastasis were significantly associated with lack of response to ICI (χ^2^-test p-value =0.006 and p-value =0.001, respectively). Other characteristics such as sex, age, or smoker status were not significantly associated with the response (Suppl. Table 1).

### Analysis of mutational landscape

We obtained a high-resolution map of the most common somatic mutations in 121 previously defined BLCA-associated genes (Figure 1, Suppl. Table 2). *TP53* was the most frequently mutated with 132 mutations followed by *KMT2D* (112 muts), *ARID1A* (78 muts), *ELF3* (67 muts), *KDM6A* (56 muts), and *FGFR3* (55 muts). These genes have also been found to be frequently mutated in BLCA tumors from TCGA ^10^ and other cancer cohorts ^20^.

We tested if there was any association between mutations in these genes and the response to the treatment. Although mutations in the *FGFR3* are a hallmark for reduced immune infiltration ^10,11^, we observed no relationship between *FGFR3* alterations and response to ICI, strengthening the conclusions of a previous report that was based on 103 patients, 17 of which had *FGFR3* alterations ^19^. The only gene that was significantly associated with response was *ARHGEF12* (mutated in 11 responders and 1 non-responder, adjusted p-value = 0.015). This gene encodes a Rho GTPase and its deletion has been associated with neuroblastoma differentiation and decreased stemness-related gene expression ^21^.

### Missense and non-stop mutations are significantly associated with the response to ICI

We measured the tumor mutational burden (TMB), which is the number of non-synonymous mutations generated in the tumor, using WES data. For patients of the UNC-108 and UC-GENOME cohorts, we used the provided TMB values, which were derived from panel DNA data. To integrate the results of the different cohorts we transformed the original TMB values to Z-scores, separately for each cohort, and then combined the Z-scores of the different cohorts (n=378).

We found that responders had significantly higher TMB than non-responders (Figure 2A and 2B), consistent with previous findings ^3,9,15,22^. While both subclonal and clonal mutations were significantly associated with the response (Figure 2C), the first ones had a larger effect size. The odds ratio obtained from a logistic regression was higher for subclonal TMB (OR = 1.55, 95%CI [1.30; 1.92]) than for clonal TMB (OR = 1.24; 95%CI [1.10; 1.42]). This is in contrast to previous findings based on one of the cohorts, HdM-BLCA-1 ^18^, in which clonal mutations were shown to have a larger influence on the response. We also observed a significant enrichment of APOBEC-induced mutations in the responder group (Figure 2D); this variable was moderately correlated with TMB (r=0.47, Figure 2E). The two APOBEC signatures SBS2 and SBS13 were significantly associated with the response, but the first one had a stronger effect (linear regression values B = 5.40 and p-value = 0.0048) than the second one (B = 1.91 and p-value = 0.0107).

**Figure 2.**
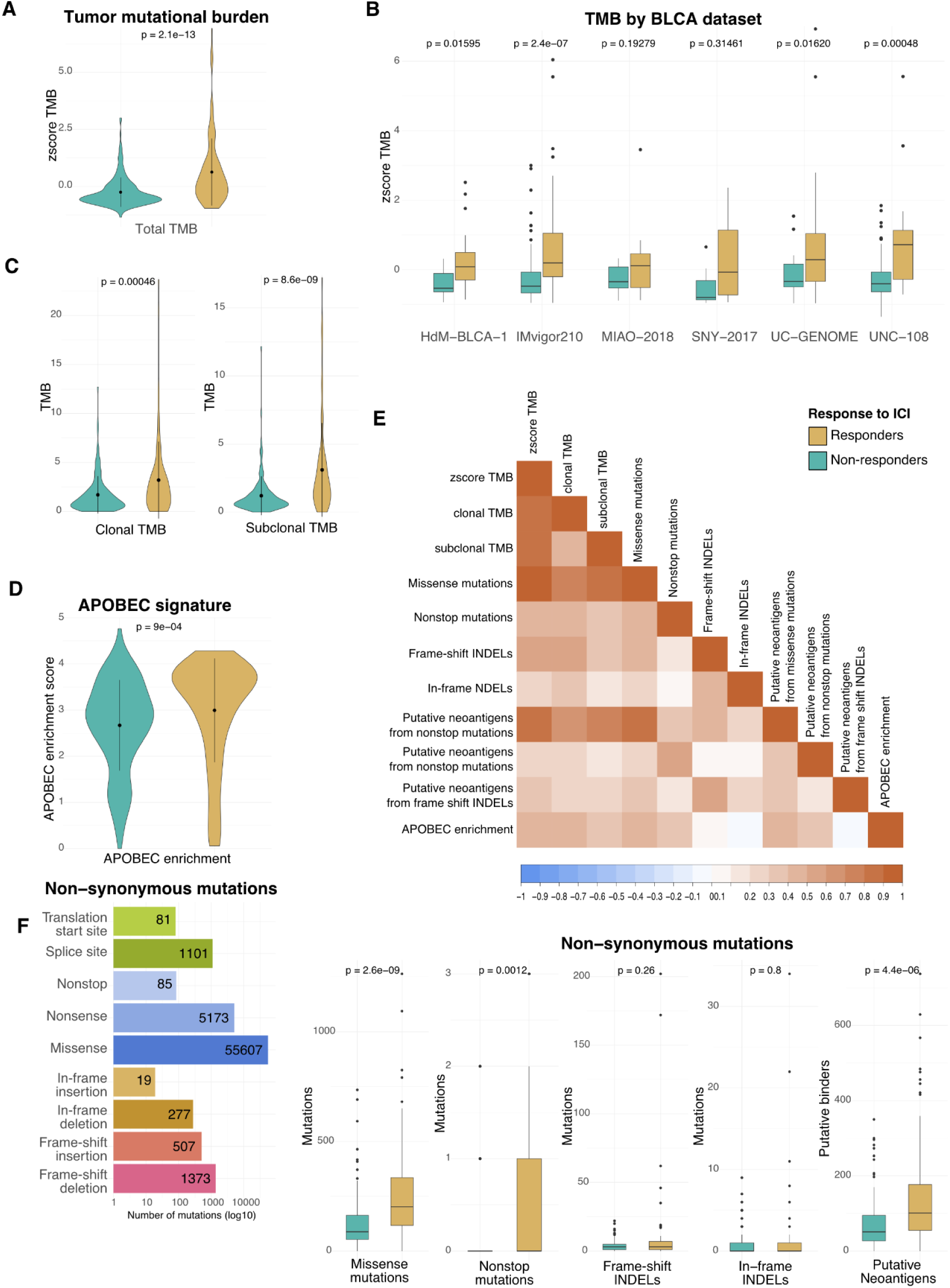
Relationship between somatic mutations and the response to ICI. **A.** Responders have a higher tumor mutational burden (Z-score TMB) than non-responders (median Z-score TMB R=0.18, NR=-0.45). TMB is calculated as the number of non-synonymous mutations per 50Mb. The differences were highly significant (Wilcoxon test, p-value < 10^-3^). **B.** Responders show a higher mean TMB than non-responders. This finding is consistent over all six datasets, and four of the six datasets display significant differences (Wilcoxon test, p-value < 0.05). **C.** The difference between treatment response groups remains when separating the TMB into clonal and subclonal by a cancer cell fraction (CCF) cutoff of 0.9. Median clonal TMB R: 2.15 mut/50Mb, NR: 1.1 mut/50Mb; median subclonal TMB R: 1.19 mut/50Mb, NR: 0.8 mut/50Mb. **D.** Responders are significantly enriched in APOBEC-induced mutations compared to non-responders (Wilcoxon test, p-value=9e-04; median APOBEC enrichment score R: 3.45, NR: 2.85). **E.** Spearman correlation between different DNA-derived variables. **F.** Number of non-synonymous mutations by type and association of the different mutation types with response. Missense mutations, nonstop mutations, and putative neoantigens were found to be significantly associated with the response (Wilcoxon test, p-value < 0.05). Medians for number of missense mutations R: 202 muts, NR: 88 muts; nonstop mutations R: 0 muts, NR: 0 muts, frameshift insertions/deletions R: 3 muts, NR: 3 muts; in-frame insertions/deletions R: 0muts, NR: 0muts; putative neoantigens R: 22.5 neoantigens, NR: 12 neoantigens. Neoantigens were predicted from missense mutations applying a threshold of 500nM IC_50_ binding affinity in NetMHCpan 4.0. P-values obtained by two-sample Wilcoxon test. R: responders, NR: non-responders, TMB: tumor mutational burden, INDELs: insertions and deletions. Number of patient samples: 234. Additional data is available from Suppl. file 2.

Missense mutations were significantly associated with response (Figure 2F, Suppl. Table 3), independently of the predicted impact of the mutation on protein function (Suppl. Figure 1). We found that non-stop mutations (single nucleotide variants that cause a stop codon loss) were significantly associated with response, but this was not the case for frameshift or in-frame insertions and deletions (INDELs). We also predicted neoantigens derived from missense mutations using the NetMHCpan software ^23^. As previously described for different cancer types ^6,18,24^, we found that the number of predicted neoantigens derived from missense mutations was significantly associated with response. Finally, we constructed peptides resulting from frameshift INDELs and the protein extensions generated by nonstop mutations. For these mutations we found no difference in the number of putative neoantigens between responders and non-responders (Suppl. Figure 1).

Somatic copy number alterations are common events in tumor cells of BLCA patients ^10^. Previous studies have suggested a positive relation between focal amplification of cyclin D1 (*CCND1*) and ICI response ^15^ and a negative association between genomic alterations in *CDKN2B* and *CDKN2A* and overall survival in BLCA patients ^25,26^. While we found *CDKN2A*/*CDKN2B* to be the most common deletion in our data (16 responders and 30 non-responders), and every second patient to have an amplification in the *CCND1* gene region (51.30%, 75 non-responders and 43 responders), none of the copy number alterations were found to be significantly associated to ICI response (Chi-square test p-value > 0.05).

### Gene expression biomarkers associated with ICI response

We used the RNA-Seq data to compute individual gene expression values, immune cell composition in the tumor sample with the CIBERSORTx algorithm ^27^, and previously defined gene signatures. As expected, immune invasion and activation biomarkers were consistently more elevated in responders than in non-responders. This included chemokines such as *CXCL9, CXCL10, CXCL11,* and *CXLC13,* the interferon-γ and granzyme A and B, and CD8A (Figure 3A and Suppl. Figure 1). We observed significantly higher expression of the immune checkpoints PD-1 and PD-L1 in responders than in non-responders (Figure 3B). While high expression of these genes is generally associated with poorer prognosis in cancer, they have also been suggested as biomarkers of ICI response ^28–31^.

**Figure 3.**
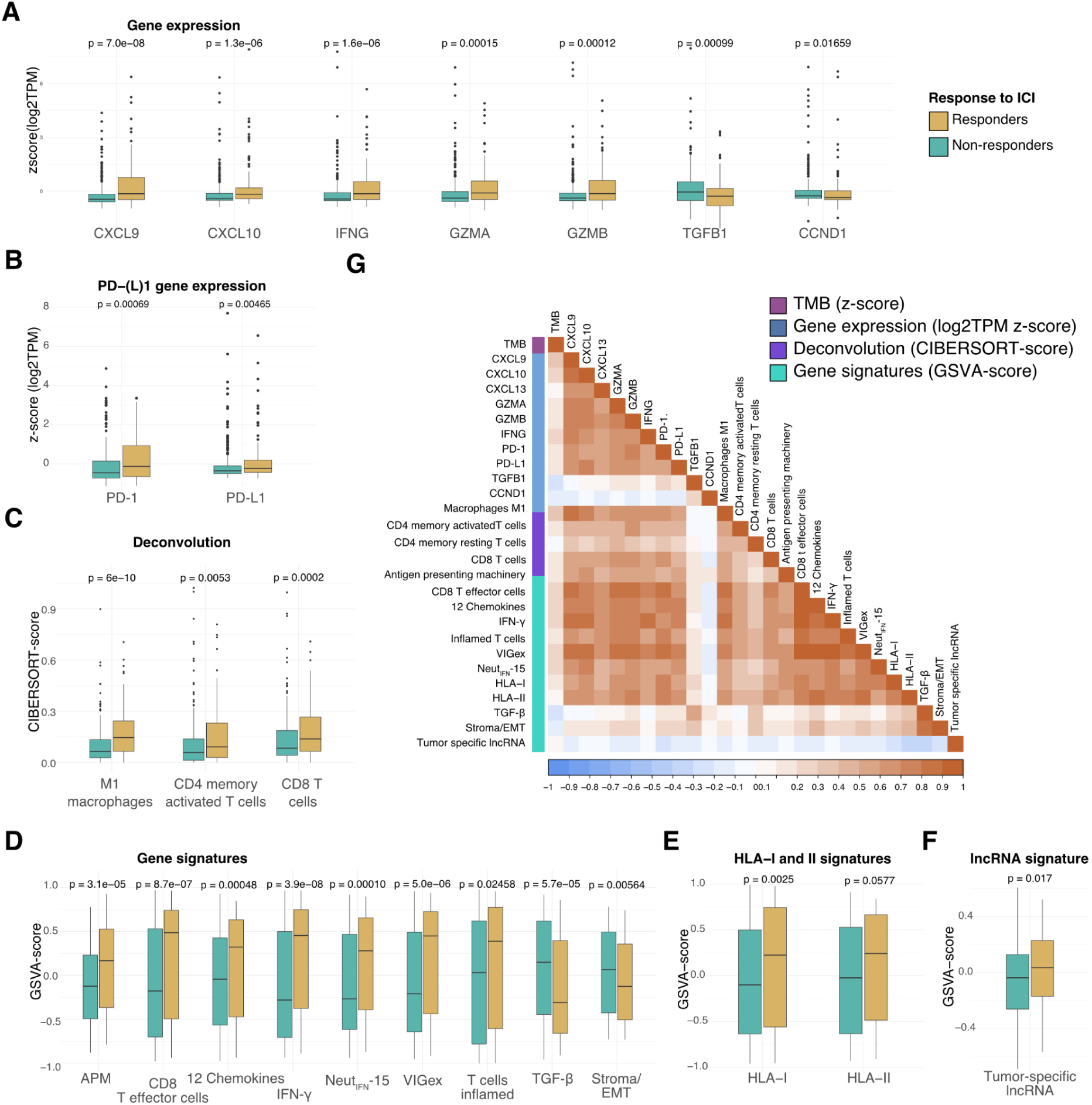
Dissecting the effects of different immune-related variables. **A**. Normalized expression values for selected genes described in the context of Immune activity or suppression (N=420). All comparisons were significant after multiple testing adjustment, except for the genes TGFB1 and CCND1. **B.** Expression of the immune checkpoint molecules PD-L1 (gene CD274) and PD-1 (gene PDCD1) is significantly higher in responders. **C.** Deconvolution analysis using CIBERSORT shows higher immune cell abundance in responders. **D.** Six signatures of tumor antigen presentation and immune response are enriched among responders, while immune suppression signature scores are higher in non-responders. All comparisons were significant after multiple testing adjustment, except T cells inflamed and Stroma/EMT. **E.** Gene signatures combining the expression of HLA-I and HLA-II types are higher in responders than in non-responders. **F.** Responders show higher expression of a signature related to tumor-specific long non-coding RNAs (lncRNA) than non-responders. **G.** Correlation matrix including TMB and RNA-Seq variables. P-values obtained by two-sample Wilcoxon test. R: responders, NR: non-responders, APM: antigen-presenting machinery, lncRNA: long non-coding RNA. Additional data is available from Suppl. file 2.

The abundance of CD8+ T cells, memory-activated CD4+ T cells and pro-inflammatory macrophages M1 was significantly higher in responders than in non-responders (Figure 3C). Three out of four gene expression signatures related to CD8+ T cell activation were significantly associated with response: CD8 T-effector from Mariathasan *et al.* 2018 for bladder cancer ^9^, CD8 T-effector from McDermott *et al.* 2018 for kidney cancer ^32^ and CD8 T-effector from POPLAR for lung cancer ^33^ (Figure 3D). CD8 T-effector from Bindea *et al.* 2013 for colorectal cancer ^34^ did not achieve statistical significance. Similarly to the gene expression of *IFNG*, responder samples were significantly enriched in both signatures of the IFN-γ pathway by Ayers *et al.* (2017)^35^ and Cristescu *et al.* (2018)^36^. The recently published gene signature VIGex, which combines different genes involved in immune response ^37^, and Neut-IFN-15, relating IFN-γ stimulated neutrophil to improved response to ICI ^38^, were also significantly associated with response (Figure 3D). We observed higher scores for the antigen-presenting machinery (APM) gene signature by Thompson et al. (2020)^39^, as well as for HLA-I, in responders than in non-responders (Figure 3E and Suppl. Figure 2).

We also investigated if tumor-restricted lncRNAs, which might contain non-canonical ORFs with the potential to generate HLA-bound peptides ^40,41^, were more prevalent in responders than non-responders. If so, this would support the hypothesis that non-coding sequences might be a source of immunogenic antigens ^42^. In each cohort, we generated a gene expression signature based on tumor-specific lncRNAs containing open reading frames with cancer-derived immunopeptidomics evidence (Suppl. Figure 3). We found that, when the signatures from different datasets were combined together, responders had significantly higher values of this lncRNA signature than non-responders (Figure 3F). When the analysis was performed separately for each cohort, only IMvigor210 achieved statistical significance (Suppl. Figure 4).

We found significantly lower expression of the immunosuppressive transcription growth factor beta 1 (TGF-β), and of the cyclin gene *CCND1,* in responders than in non-responders (Figure 3A). A biological pathway supporting tumor progression is the epithelial to mesenchymal transformation in stroma cells (Stroma/EMT)^9,43^ was also negatively associated with response (Figure 3D).

### Relationship between different signatures

The different immune activation signatures tested were all strongly positively correlated with each other (Figure 3G, Suppl. file 2). Intriguingly, the immunosuppressive stroma/EMT signature, which was weakly positively correlated with immune activating signatures, showed a negative correlation with TMB (r=-0.169, p-value = 0.002). *CCND1*, more abundant in non-responsive tumors, was positively correlated with TGF-β (r=0.177, p-value < 0.001). PD-1 and PD-L1 expression levels, as well as HLA-I and HLA-II signatures, showed a positive correlation with CD8+ and CD4+ activating cells (r=0.4-0.65, p-value<0.001). The lncRNA signature showed no correlation with the other biomarkers studied. Principal component analysis identified five major groups of variables that contribute to explain the variability across tumor samples: 1. Mutation status (TMB/APOBEC), 2. Immune activation signatures (CD8+ T cells, IFN-γ, etc.), 3. TGF-β/EMT immuno-suppressor signatures; 4. tumor-specific lncRNAs and, 5. CCDN1 (Suppl. Figure 5).

### The determinants of the response to ICI response depend on the subtype

The analysis of BLCA samples from the TCGA project identified five subtypes based on different biomarkers ^10^. We used this well-established methodology to classify the tumors in our meta-cohort. The largest group of patients was luminal infiltrated (52 R & 116 NR), followed by basal-squamous (35 R & 78 NR), luminal-papillary (31 R & 56 NR) and luminal (11 R & 25 NR) (Figure 4A). Neuronal represents the smallest subtype (10 R & 6 NR). The only subtype with a significantly better response to treatment than the general trend was neuronal (62.5% *versus* 33% of responders, Fisher exact test, p-value = 0.014).

**Figure 4.**
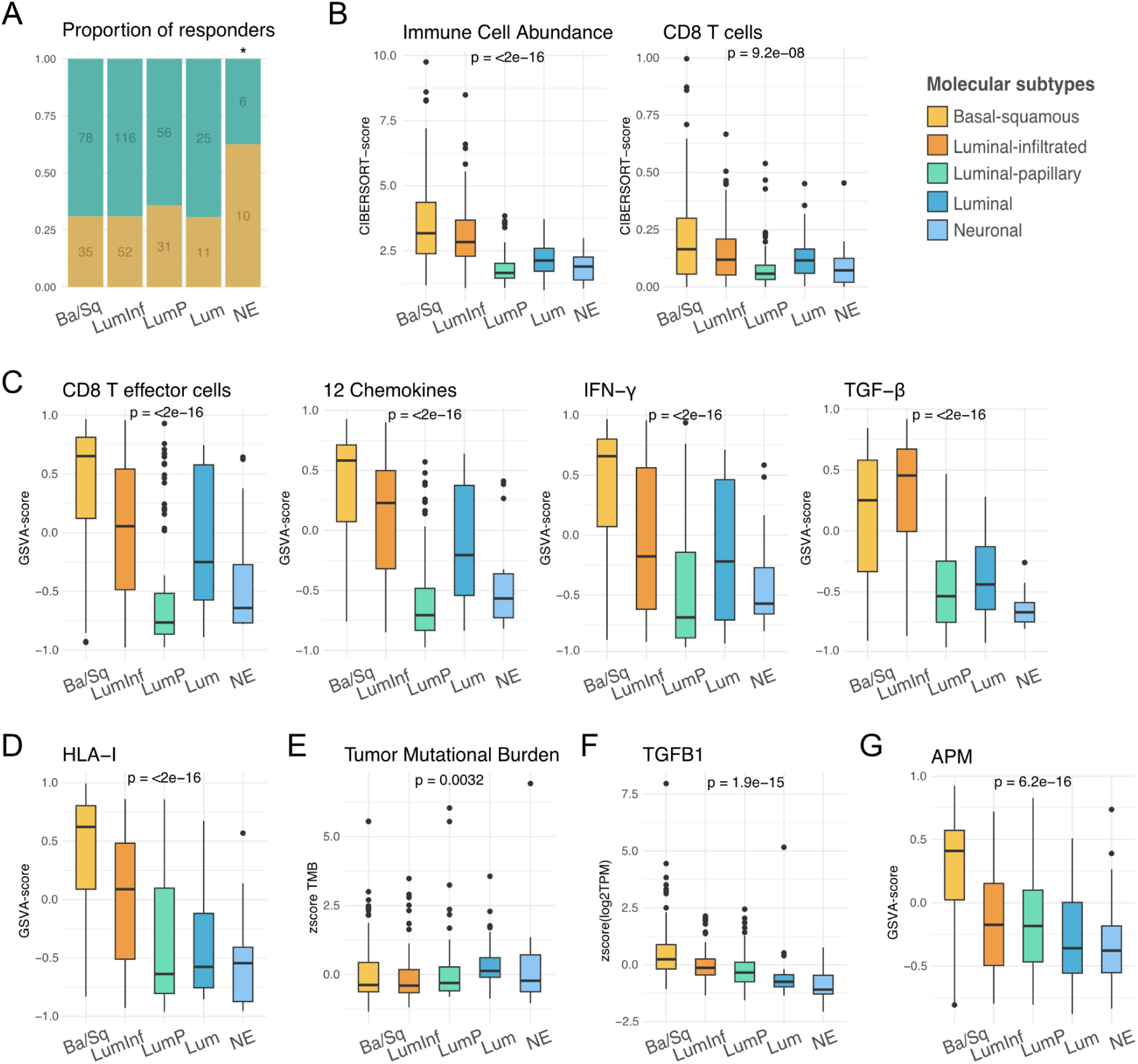
The TCGA-subtypes luminal and neuronal have higher TMB, while basal-squamous and luminal-infiltrated show higher immune activity. **A.** Proportion of responders and non-responders in each of the five TCGA subtypes. TMB was weakly correlated with the immune activation signatures. The neuronal subtype is the only one with a significant excess of responders over non-responders (Fisher exact test, p-value = 0.014, indicated with *) **B.** Basal-squamous and luminal-infiltrated have high values of immune cell abundance (absolute score obtained from CIBERSORT), while luminal, luminal-papillary and neuronal show low immune invasion. Immune cell abundance of CD4 T cells is similarly high in basal-squamous and luminal-infiltrated subtypes (CIBERSORT). Luminal-infiltrated shows the highest mean of CD4 memory resting T cells (CIBERSORT). **C.** Basal-squamous and luminal-infiltrated have the highest expression of immune biomarkers, both immune activating and suppressive markers. **D.** HLA expression is also highest in luminal-infiltrated and basal-squamous. **E.** TMB is highest in the luminal and neuronal subtypes. **F.** The tumor-infiltrated subtypes basal-squamous and luminal-infiltrated show the highest values of TGF-β gene expression. **G.** Basal-squamous is enriched in antigen-presenting machinery (APM) compared to other subtypes. N=420. Overall p-values for the inter-group comparison were obtained by Kruskal-Wallis t-test. R: responders, NR: non-responders. Additional data is available from Suppl. file 2.

We observed that 30% of luminal-papillary tumors were *FGFR3*-altered (26 out of 87), followed by 8% for luminal. Basal-squamous and luminal-infiltrated had high immune cell infiltration, as expected^10^ (Figure 4B and 4C, respectively; Suppl. Figure 6). In addition, we observed that these two subtypes had higher HLA I gene expression (Figure 4D) and, at least for basal-squamous, higher expression levels of the antigen presentation pathway (Figure 4G). They also had high TGF-β and low TMB values (Figures 4D-4F).

We next analyzed the different variables in the immune-infiltrated (luminal infiltrated and basal-squamous) and non-immune-infiltrated (luminal-papillary, luminal and neuronal) subtypes separately. Although immune-infiltrated tumors had significantly lower TMB than non-immune infiltrated ones (Figure 5A), TMB and APOBEC enrichment were significantly associated with response (Figure 5B). The effect of immune activation and immunosuppression signatures on the response was significant in both groups, but in general more marked in the case of the immune-infiltrated subtypes (Figure 5C and 5D, Suppl. Figures 7 and 8). For the IFN-γ signature, the difference between the median of the response group and the median of the non-response group was 0.727 for immune-infiltrated and 0.229 for non-immune-infiltrated. We observed the same trend for the CIBERSORT score (0.364 and 0.112, respectively), M1 macrophages (0.099 and 0.037, respectively) and CD8 T cells (0.07 and 0.026, respectively). For lncRNAs, the signature was only significant for the immune-infiltrated group (Figure 5E).

**Figure 5.**
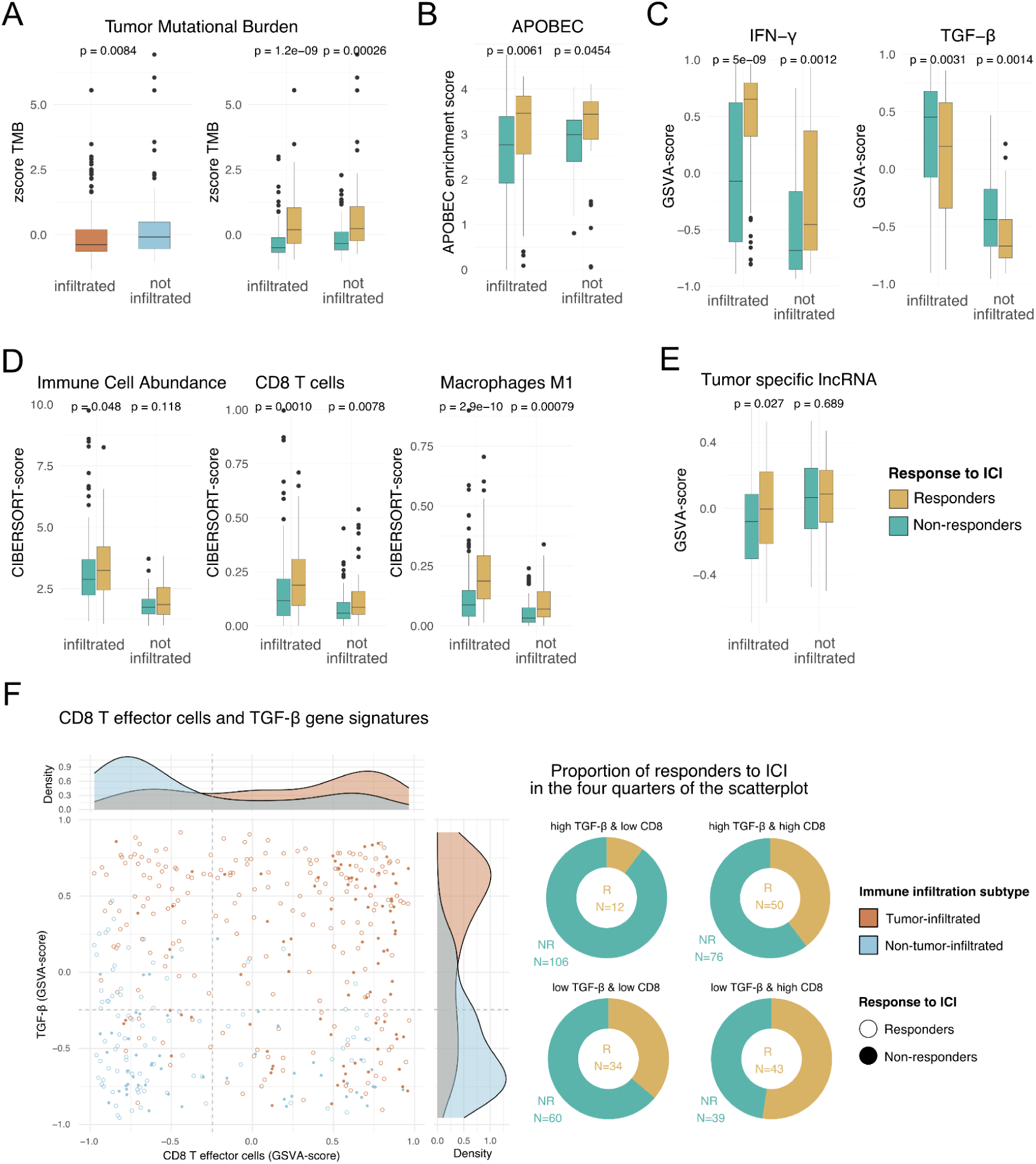
Influence of different biomarkers in the immune-infiltrated and non-immune-infiltrated subtypes. **A.** Overall, TMB is higher in the non-immune infiltrated group. In both groups, responders have significantly higher TMB. **B.** In both groups, responders have higher enrichment scores of the APOBEC mutational signature. **C.** Responders have higher IFN-γ expression values, and lower TGF-β expression values than non-responders for both subgroups. **C.** CIBERSORT score is higher in responders of the infiltrated groups, but not the non-infiltrated group. Differences between responders and non-responders are significant for CD8+ T cells and macrophages M1. **D.** Only in the infiltrated group, responders have higher expression of the lncRNA signature. No difference was observed between response groups in the non-immune-infiltrative group. P-values obtained by two-sample Wilcoxon test: R: responders, NR: non-responders. **F.** Relationship between CD8 T cell and TGF-β gene expression. Immune-infiltrated samples tend to have high CD8+ T cell abundance, and in many cases also high levels of the TGF-β signature. Dashed line marks the optimal cutoff for each gene signature obtained by ROC (TGF-β: 0.0163; CD8 t effector cells: 0.246). Additional data is available from Suppl. file 2.

Because of the high TGF-β expression values detected in the immune-infiltrated types (Figure 5C), we decided to investigate in more detail the relationship between immune activation and immune suppression signatures. We found a relatively large group of patients with high CD8+ T cell infiltration and also high TGF-β values, these patients had a lower probability to respond than those with high CD8+ and low TGF-β values (39.7% *versus* 52.4% of responders, Fisher exact test p-value = 0.0479, Figure 5F). When we performed the analysis per subtype, the same trend was observed, especially for luminal infiltrated, although the results did not achieve statistical significance (Suppl. Figure 9). These observations can explain why tumors with high immune infiltration respond less well to the treatment than expected.

### Predictive models integrating different omics variables

Based on the above-described findings, we selected a list of representative variables to construct predictive models of the response to ICI. The complete set of variables comprised mutation-based variables (TMB, non-stop mutations, APOBEC-enrichment score), gene expression of selected genes (*PD-1, PD-L1, CCND1*), immune signatures (IFN-γ, stroma/EMT, inflamed T-cells, TGF-β and antigen-presenting machinery pathways), a signature of tumor-specific lncRNAs, immune cell abundance (M1 macrophages, CD4 memory activated T cells, CD8 T cells and regulatory T cells), as well as clinical information (ECOG, liver metastasis).

A random forest model trained with these variables achieved high accuracy, as indicated by an area under the curve (AUC) of 0.761 (Figure 6A, Suppl. Table 4). In clinical practice, the number of mutations in a tumor sample is a widely used marker for decision-making in the context of ICI therapy ^44^. A random forest model considering only z-score TMB had an AUC of 0.678. A threshold model in which tumor samples with more than 10 mutations/Mb were predicted to be from responders and those with less mutations from non-responders was associated with an AUC of 0.61 (Suppl. Figure 10).

**Figure 6.**
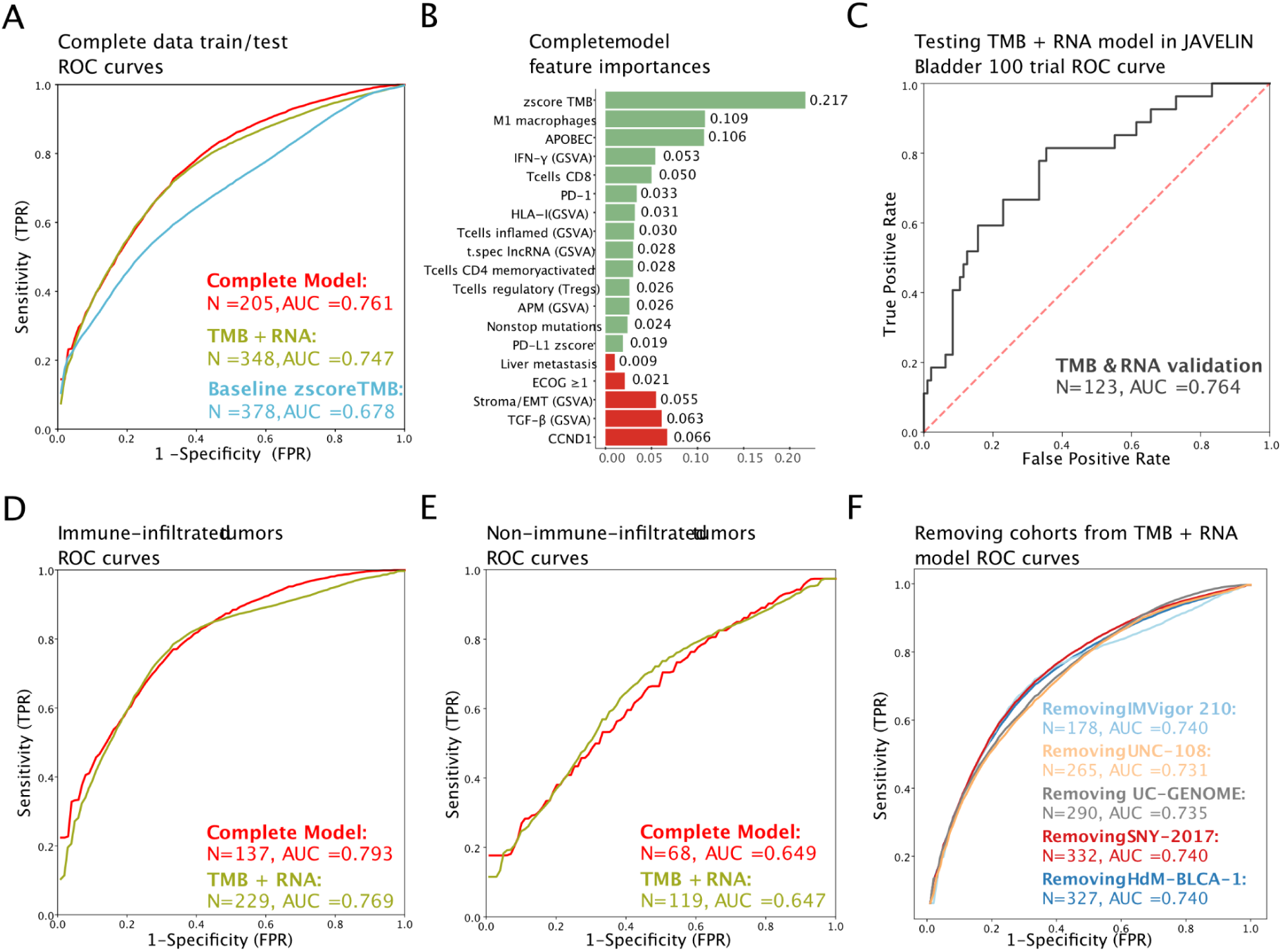
Predictive models of the response to immunotherapy. **A.** ROC curves for the complete model, TMB and TMB+RNA (TMB and RNA-seq derived variables), with AUC and number of samples (N). FPR: false positive rate; TPR: true positive rate. The average of 1000 runs is shown. **B.** Feature positively or negatively associated with response. Length of the bar represents feature importance from random forest. Color reflects association with response taken from the previous manuscript sections (green: positively associated, red: negatively associated). **C**. ROC curve for testing the TMB+RNA model in the validation cohort JAVELIN Bladder 100 trial. AUC and N values are indicated. **D.** ROC curve of the complete model and TMB+RNA for the subset immune-infiltrated tumors, with AUC and N values. **E.** ROC curve of the complete model and TMB+RNA for the subset of non-immune-infiltrated tumors, with AUC and N values. **F.** ROC curves for TMB+RNA model when removing one of the cohorts at a time.

Random forest models provide information on the contribution of each variable to the model (feature importance). The variables with a clear association with response in the complete model were, in descending order, TMB, M1 macrophages, APOBEC-enrichment, IFN-γ signature, CD8+ T cell abundance, PD1 gene expression and HLA I signature (feature importance > 0.03, Figure 6B). Those associated with lack of response were, also in descending order, CCND1 gene expression, Stroma/EMT signature and TGF-β gene signature.

As a validation cohort, we tested the model on data from the JAVELIN Bladder 100 trial ^45^, which includes advanced urothelial cancer patients treated with avelumab (anti-PD-L1). While no raw data was available, we were provided with processed data. With this data, we tested the model based TMB + RNA using data from 123 patients from the JAVELIN Bladder 100 trial (27 responders, 96 non-responders). The model achieved an AUC of 0.764 in the validation run, surpassing the averaged AUC of 0.747 obtained from 1000 seeds in the train/test phase (Figure 6C). This result underscores the robustness of our model, highlighting its potential in predicting outcomes for patients treated with immune checkpoint inhibitors.

We then built models separately for the immune-infiltrated and the non-immune-infiltrated classes. Maximum accuracy was achieved in the immune-infiltrated group using all the variables (Figure 6D, AUC=0.793). The IFN-γ signature, as well as *CD8* gene expression, gained weight in this model with respect to the model developed for all patients (Suppl. Figure 11). *CCDN1* abundance was the most negative factor, followed by TGF-β levels. In contrast, the non-immune-infiltrated model showed relatively low accuracy (Figure 6E, AUC=0.649). The model based on RNA-Seq data plus TMB had an AUC of 0.769 for immune-infiltrated (n=229) and 0.647 for non-immune-infiltrated (n=119), similar to the complete model.

The robustness of the model was further tested by removing one dataset at a time. For this, we employed the model constructed with TMB and RNA-Seq data, as it included more patients than the complete model (348 vs 205) and showed almost the same accuracy (AUC 0.747 vs AUC 0.761, Figure 6A). Removing individual datasets resulted in models of similar accuracy (Figure 6F), indicating that the model does not suffer from over-fitting. The fact that the number of non-responders was approximately double than the number of responders did not seem to have a significant effect either (Suppl. Figure 12).

To gain further insights into the drivers of the response to ICI in the non-immune-infiltrated samples, we derived subtype-specific decision trees and inspected the distribution of the variables across patients (Suppl. Figures 13 and 14). In the case of luminal papillary, the largest group, we also built a random forest model (n=71; Suppl. Table 5). This model identified PD-1 z-score as the third most important feature following macrophages M1 and TMB. The decision tree also indicated that high PD-1 expression was strongly associated with response, with 11 out of 14 patients with PD-1 z-score >-0.41 being responders. Among the remaining 15 responders, 13 had very low values of the stroma/EMT gene signature (stroma/EMT <-0.527). Therefore, a large part of the responders, 24 out of 26, were characterized by either high PD-1 values or low stroma/EMT.

In the case of luminal, the decision tree that we obtained revealed an association between absence of Regulatory T cells (Treg) and response to ICI. The percentage of responders with Treg values above 0.003 was only 9% (1 out of 11), whereas it was 58% for non-responders (14 out of 24)(p-value = 0.0095). Regulatory T cells are a subset of CD4^+^ T cells with immunosuppressive effects ^46^. This effect was only observed in the luminal subtype (Suppl. Figure 14).

In the neuronal subtype, the lack of response was invariably associated with low AMP signature values (AMP z-score <-0.36 in all non-responders, see also Suppl. Figure 8). So, the inability to present antigens at sufficiently high levels seems to be a strong determinant for the treatment to fail in this group of patients.

## DISCUSSION

We have compiled the largest clinical, mutation and gene expression dataset from patients with advanced BLCA treated with ICI. Reanalysis of previously associated signatures has confirmed that tumor mutational burden (TMB) and APOBEC-induced mutations are consistently associated with the response ^3,15,18,47,48^. Frameshifts mutations, reported to be significant in melanoma ^49^, were not found to be significant here. Analysis of RNA sequencing data has confirmed that immune activation biomarkers were significantly associated with response to ICI, as previously reported for different cancer types ^15,35,36^. Macrophage M1 and CD8+ T cell levels were higher in responders than in non-responders, even in those cancer subtypes with relatively little immune infiltration. Two factors that negatively affected the response were TGF-β/stroma/EMT and *CCND1* gene expression levels. It has been proposed that TGB-β negatively affects the response to ICI by blocking T cell penetration in the tumor ^9^. In the case of *CCND1*, a previous association had been found between *CCND1* amplification and lack of response ^15^. The inhibition of the protein complex cyclin D1/CDK4 results in higher PD-L1 levels, linking the kinase activity modulated by *CCND1* with cancer immune surveillance ^50^. Our results indicated that, in least in bladder cancer, measuring *CCND1* gene expression could have a stronger prognostic value than measuring copy number.

We found several new biomarkers of the response to ICI. The first one was the number of non-stop mutations, which was positively associated with response. Although non-stop mutations are only present in about 21% of the tumors, the large number of patients analyzed here allowed us to draw this association. Non-stop mutations can extend the coding sequence and, as a result, “foreign” antigens might be formed. Another signature was linked to the tumor-specific activation of lncRNAs, which are an alternative source of tumor-specific antigens ^42,51^. Mutations in the gene ARHGEF12 were also significantly associated with response. This gene encodes a yet poorly characterized Rho GTPase. Interestingly, a study comparing pre-and posttreatment samples of two cancer patients treated with anti-PD-1 reported that tumor clones with *ARHGEF12* mutations were only detected in pretreatment samples ^52^. This could imply that anti-PD-1 treatment was effective in eliminating cells containing mutated *ARHGEF12*.

Previous models of the response to immunotherapy included data from different types of cancer ^15, 17,53,54^ or focused on specific BLCA cohorts ^16^. The size of our study allowed us to investigate the determinants of the response to ICI in different BLCA subtypes. Surprisingly, we found that the two subtypes with the highest immune infiltration-basal squamous and luminal infiltrated-did not show an overall better response than the other subtypes. This is likely to be relevant for trials that select treatment on the basis of cancer subtype ^55^. We discovered that the immune-infiltrated subtypes tend to have lower TMB values, as well as higher immunosuppressive signatures, than other subtypes, and these factors decrease the likelihood of responding. For immune-infiltrated subtypes we achieved high accuracy in response prediction models that included RNA-Seq data in addition to TMB. This encourages the use of expression biomarkers, such as signatures of macrophages M1, CD8+ T cells and TGF-β, in predicting the outcome of the treatment for these cases. For non-immune-infiltrated subtypes, other biomarkers appeared to be particularly relevant. PD-1 expression levels were strongly associated with response in the case of luminal papillary and basal-squamous, but were of little relevance in the other subtypes. The potential inhibitory effect of Treg cells was consistent with the larger number of these cells in the tumors of non-responders versus non-responders, in the case of the luminary subtype.

Anti-PD-(L)1 therapy depends on the expression of PD-1 and PD-L1, which can be quantified using immunocytochemistry prior to treatment ^2,56^. The expression of these genes was significantly associated with response. However, it was also more elevated in the immune-infiltrated tumors than in the non-immune-infiltrated tumors, independently of the response. This illustrates how taking into account the specific tumor subtype, or the degree of immune inflammation of the tumor, might be key for a correct interpretation of these biomarkers in a clinical setting.

Although a large number of variables were considered here, there may be yet unknown factors that also have an influence in the response. The neuronal subtype showed a strikingly high response rate. It has been hypothesized that this kind of tumor could be more responsive to ICI due to the expression of tissue-restricted neuronal or neuro-endocrine proteins, together with low TGF-β gene expression values ^14^. The present study, which included a larger number of patients, confirmed that TFG-β levels are particularly low in this class. As already mentioned, the non-responders (6 out of 16) were all characterized by low values of the antigen-presentation machinery. This reinforces the fundamental role of tumor-specific antigens, of mutation origin or otherwise, in mediating the response to the treatment.

Using part of the data for validation we obtained an AUC estimate of 0.761 for the model that used the complete set of variables (0.793 for the immune-infiltrated subtypes) and 0.747 for the RNA + TMB model (gene expression variables and TMB). This represented an improvement over using TMB alone (0.678). When we tested the RNA + TMB model against the JAVELIN Bladder 100 trial we obtained an AUC of 0.764. These predictive values are in the range of those obtained by a previously obtained model that combined data from different types of cancer ^15^; the AUC values were between 0.66 and 0.86 depending on the test dataset. The elastic net logistic regression model by Damrauer et al. (2022)^16^, built on the bladder cancer IMvigor210 dataset, showed an AUC of 0.84 when tested on the validation portion of the same dataset, 0.82 when tested on UNC-108 and 0.65 when tested on UC-GENOME.

In this study we integrated disparate cohorts of bladder cancer patients, and this poses a number of challenges. In order to control for sequencing batch effects we transformed the data to normalized z-scores, which allowed us to merge datasets. Not all the datasets included the same set of variables, therefore, the complete model did not include all patients with ICI response information. We tested the effect of leaving out one cohort of a time, or employing a similar number of non-responders than responders. External cohort validation using the independent JAVELIN Bladder 100 cohort demonstrated the robustness of our model. In this specific cohort the patients had been treated with chemotherapy prior to immunotherapy, but this does not seem to impact the results. Testing the model in cohorts treated with anti-PD-(L)1 in combination with other drugs (e.g. Enfortumab vedotin) can be foreseen.

In conclusion, we have combined six datasets to build the largest bladder cancer-specific meta-cohort of ICI therapy, and conducted a comprehensive multi-omics analysis. The results have uncovered new ICI associated variables and have shed light on the complex interplay between tumor biology, the immune microenvironment and treatment response. Our study underlines the importance of subtype-specific factors for more personalized treatment strategies and enhanced patient outcomes in the era of immunotherapy.

## METHODS

### Omics datasets

We accessed tumor RNA-Seq, paired tumor-germline WES and targeted DNA sequencing from six advanced or metastatic bladder cancer cohorts. All patients received anti-PD-1/PD-L1 treatment. Germline WES was conducted on whole blood samples, tumor WES and RNA-Seq data obtained from FFPE samples coming from primary or metastatic cancer tissue.

1. IMvigor210, N=348, (Mariathasan et al., 2018)
EGA study ID EGAS00001002556
[https://ega-archive.org/studies/EGAS00001004343]
2. HdM-BLCA-1, N=27, Boll et al., (2023)
EGA study ID EGAS00001007086
[https://ega-archive.org/studies/EGAS00001007086]
3. SNY-2017, N=25, Snyder et al. (2017)
dbGaP accession ID phs001743
[https://www.ncbi.nlm.nih.gov/projects/gap/cgi-bin/study.cgi?study_id=phs001743.v1.p1]
4. MIAO–2018, N=26, Miao et al. (2018)
dbGaP accession ID phs001565
[https://www.ncbi.nlm.nih.gov/projects/gap/cgi-bin/study.cgi?study_id=phs001565.v1.p1] and phs000694
[https://www.ncbi.nlm.nih.gov/projects/gap/cgi-bin/study.cgi?study_id=phs000694.v3.p2]
5. UC-GENOME, N=191, Rose et al. (2021)
dbGaP accession ID phs003066
[https://www.ncbi.nlm.nih.gov/projects/gap/cgi-bin/study.cgi?study_id=phs003066.v1.p1]
6. UNC-108, N=89, Damrauer et al. (2022)
Omnibus accession ID GSE176307
[https://www.ncbi.nlm.nih.gov/geo/query/acc.cgi?acc=GSE176307]

Three patient samples were excluded due to corrupted raw data files (IMvigor210: 10059, 10119, SNY-2017: 9881). Patient 4072 of the SNY-2017 cohort was excluded due to low tumor mean coverage according to the publication.

As a validation cohort, data of the JAVELIN Bladder 100 cohort (ClinicalTrials.gov identifier: NCT02603432) was obtained from Merck Healthcare KGaA under a data sharing agreement. The dataset is derived from biological samples of advanced bladder cancer patients, randomly assigned to chemotherapy best supportive care alone or in combination with the anti-PD-1 drug avelumab. From this cohort, we included 123 patients ranked as responder or non-responder to immunotherapy following RECIST1.1 criteria.

### Single nucleotide variant calling

All paired tumor and germline WES samples were processed using the same pipeline following the GATK pipeline (version 4.0.1.2). Raw sequencing reads were trimmed using cutadapt (version 4.1), the quality was controlled using fastQC (version 0.11.7). Reads were aligned to the reference genome GRCh38 using BWA (version 0.7.17), and, in cases of several fastq files per patient, merged into one BAM file using GATK MergeSamFiles. Next, intervals and INDELs were realigned using the GATK RealignerTargetCreator and IndelRealigner to finally calibrate base scores and mark duplicates using GATK BaseRecalibrator. A summary file of the final BAM file was built using GATK GetPileupSummaries, contamination was estimated with GATK CalculateContamination and coverage using qualimap (version 2.2.1). A metrics file was built using GATK CollectAlignmentSummaryMetrics. Mutations were called with the three callers GATK Mutect2, Strelka2 (version 2.9.10) and VarScan2 (version 2.4.4). We used SAMtools (version 1.12) mpileup to build the input for VarScan2. Additionally to the default filters, Mutect2 was run using the germline-resource file somatic-hg38_af-only-gnomad.hg38.vcf.gz from gatk hg38 and GATK FilterMutectCalls using the contamination file previously generated. For all callers only mutations flagged as PASS were kept. We then build an ensemble file of mutations detected by a minimum of two out of three callers using bcbio-variation ensemble (version 0.2.2.6) and annotate them with VEP (version 104). Maf files were generated using vcf2maf (version 0.1.16). Final filters applied to mutations were a population-wide allele frequency under 5% (gnomAD), a minimum sample depth of 30, and a minimum alternative allele depth of 3. Maftools R-package (version 2.10.0) was used for TMB calculation (non-synonymous mutations per capture size of 50Mb)^57^. For two patients of the IMvigor210 cohort, the variant callers detected zero somatic mutations (10124, 10299). In patient 6428 of the SNY-2017 dataset, none of the called mutations passed the filter. We identified a total of 64,275 somatic non-synonymous mutations in 15,439 genes. This represents an average of 202 non-synonymous mutations per patient (318 patients with WES data). If we only considered the 236 patients classified as responders or non-responders and with WES data, the average was similar (210 mutations per patient).

The selection of tested tumor suppressor and oncogenes was obtained from the following sources: COSMIC (https://cancer.sanger.ac.uk/cosmic/census?tier=2, last accessed September 11th 2023) as well as reported chromatin modifying genes, chromatin regulating genes and common bladder cancer genes. A complete list is provided in Suppl. file 2.

### Somatic copy number alterations

The ASCAT R-package (version 3) was used to obtain tumor purity and local allele-specific copy number estimates from WES samples. ASCAT was run following the recommendations of version3 for WES data. AlleleCount (version 4.3.0) was used to generate LogR and BAF files. The necessary loci file was downloaded from the nf-core/sarek pipeline (release 3.1.) and subsetted for loci of the according BED file. We have tested differences between responders and non-responders using Chi-square tests for somatic copy number alterations of a selection of genes significantly amplified or deleted in TCGA (taken from Suppl. table 2 of Zack et al., 2013)^58^.

### Mutation clonality

Mutations with a cancer cell fraction (CCF) above 0.9 were considered clonal. CCF was calculated as: CCF = VAF/p * (2* (1-p) + c*p) with c being the copy number at the mutation position and p the tumor sample purity ^18^. Patients with a tumor sample purity of 1 were excluded from the clonality analysis.

### APOBEC enrichment estimation

To evaluate the enrichment of the apolipoprotein B mRNA editing catalytic polypeptide-like (APOBEC) mutational signature, we used the trinucleotideMatrix function from the maftools R-package (version 22) assessing the enrichment of C>T mutations within TCW motifs relative to the overall count of C>T mutations compared against a background of cytosines and TCW motifs. Samples with an enrichment score exceeding 2 and a p-value below 0.05 were deemed statistically significant. Additional mutational signatures were obtained with the deconstructSig R-package ^59^ with COSMIC signatures and normalized by exome trinucleotide context (tri.counts.method = ‘exome’).

### Neoantigen and HLA prediction

The 4-digit HLA genotype corresponding to each patient was determined using the nf-core/hlatyping pipeline (release 1.2.0)^60^ and the OptiType HLA genotyping algorithm (version 1.3.5)^61^. OptiType was executed with default parameters on blood samples to ascertain the HLA alleles of each patient. Subsequently, we generated all possible 9-mer peptides encompassing a mutation and their non-mutated counterparts. The peptide sequences were obtained from Ensembl GRCh38. Binding affinity of tumor peptides to MHC-I molecules was predicted using NetMHCpan (version 4.0)^62^ applying an IC_50_ threshold of 500 nM.

### Gene expression quantification

Total stranded RNA-Seq reads for all cohorts but UNC-108 were quality assessed using both FastQC (v0.11.5) and FastQScreen (v0.14.0) software 65. Bulk RNA-Seq sequencing reads from tumor samples were aligned to the human reference genome GRCh38 reference and the gencode annotation (version 41) using STAR (version 2.7.8)^63^. We checked the strandedness of the dataset with the RSeQC program ^64^ and, according to the result, we used the appropriate mode of featureCounts ^65^, from the Subread package (version 2.0.3), to quantify the gene expression. Normalization was performed by converting counts to transcripts per million (TPM). We integrated the information by using z-score transformation of log2(TPM) values in each of the datasets.

### Analysis of RNA-Seq data

Deconvolution values were obtained per dataset using the CIBERSORTx method on the matrices of TPM expression with the LM22 gene signature and B-mode batch correction in absolute mode ^27^. A selected list of gene signatures previously described to be associated with ICI response was tested using the Bioconductor package GSVA ^66^. GSVA scores were calculated per dataset and then merged for analysis. We computed the following gene signatures: CD8 T effector cell signature ^9^, CD8 T effector cells from McDermott ^32^, CD8 T effector cells POPLAR ^33^, 12 chemokines ^67^, IFN-γ ^35^, IFN-gamma from Cristescu ^36^, Stroma and EMT ^43^, inflamed T cells ^35^, TGF-β ^9^, VIGex ^37^, APM ^39^, NeutIFN-15 (Interferon-stimulated neutrophils) ^38^ and signatures for HLA-I and HLA-II including the protein-coding HLA genes. A complete list of the genes included in the signatures can be found in the Suppl. file 2.

The gene expression data was also used to classify the samples by molecular bladder cancer subtype BLCAsubtyping R packages (version 2.1.1)^10^.

### Data normalization

We observed that the tumor samples from IMvigor210 had higher WES sequencing coverage than the other datasets while, among germline samples, HdM-BLCA-1 samples showed the highest coverage (Suppl. Figure 15). These differences, however, did not influence the calling of mutations, as WES coverage and TMB were not correlated (Suppl. Figure 15).

The purity of the tumor samples was quite homogenous across datasets, with a mean value of 0.569 (Suppl. Figure 15). As expected, we identified a significant negative correlation between the tumor sample purity estimate obtained from ASCAT and the absolute immune cell infiltration score from CIBERTSORT (r=-0.469, p-value<0.001).

The TMB estimates from the targeted DNA sequencing datasets UC-GENOME and UNC-108 had higher values than the estimates from WES data (Suppl. Figures 16 and 17). To correct for this effect, we normalized the TMB values by dataset using z-scores.

### RECIST and clinical data

For the analysis of the association of biomarkers to immunotherapy response, we restricted the cohort to patients responding to the treatment (partial responders and complete responders) and not responding (progressive disease) following the RECIST1.1 criteria. Patients with a RECIST status of stable disease were not considered for the comparison of response groups. Although several studies have included the stable disease status as non-responders ^91516^, we aimed for a more conservative approach with the intention of a clearer separation between the response groups, as done previously by other groups ^36^. Out of the initial 707 patients, there are 56 complete, 107 partial responders and 303 patients with progressive disease status. Clinical data was collected from the corresponding publications and data repositories. The clinical variables found to be significantly different between responders and non-responders using a Wilcoxon t-test were ECOG ≥ 1 and liver metastasis. For 202 patients, no information on liver metastasis status was available and there was no ECOG evaluation for 44 patients.

### lncRNA signature

We generated a signature of tumor-specific long non-coding RNAs (lncRNA) expression (Suppl. Figure 3). The lncRNAs included transcripts annotated as intergenic long non-coding RNAs as well as processed pseudogenes. Only lncRNAs with an expression above the 75% threshold for each dataset were considered. Gene expression data from a collection of 54 tissues from The Genotype-Tissue Expression (GTEx) project was collected. LncRNAs with significant expression (median expression higher than 0.5 TPM) in any non-reproductive tissue were discarded, resulting in the generation of lists of potential tumor-specific lncRNAs per cohort.

Additionally, we selected lncRNAs with matches in the publicly available cancer-associated immunopeptidomics dataset published by Chong and co-workers ^68^. This restricted our initial list of lncRNAs to those generating peptides that could specifically bind HLA receptors in melanoma. This was done per dataset independently; a GSVA score per patient was calculated as a gene signature of the selected tumor-specific lncRNAs. The list of lncRNAs selected for each of the cohorts with RNA-Seq data can be found in the Suppl. file 2.

### Response prediction models

The machine learning framework developed for this project consists of three different components (Suppl. Figure 18). The first one comprehends all the preprocessing steps such as scaling and encoding. In the training phase, the best hyperparameters for the ML algorithm are chosen via 15-fold-cross-validation and repeating this process multiple times (100 or 1000) in order to ensure that the parameters do not fit a particular seed. Following training, the model undergoes evaluation, where two different strategies are implemented. The first one is the commonly used train/test method. In our case, we separated the data by using a 70/30 split, leaving 30% of the data unseen by the predictive model. Using the randomly selected 70% we perform a 15-fold cross validation for training the model. We tested with the remaining 30% of the data and obtained all the metrics (including AUC).

For the model we selected a set of variables that were found to be significantly associated with the response and which showed relatively low correlation among themselves and the least possible overlap in cases of gene signatures. The complete model includes the following variables: mutation-based variables (TMB (z-score muts/Mb), nonstop mutations (muts), APOBEC-enrichment score), gene expression of selected genes (z-score (log2TPM) of PD-1, PD-L1, CCND1), immune signatures (GSVA-scores of IFN-γ, stroma/EMT, inflamed T-cells, TGF-β and antigen-presenting machinery pathways) and a signature of tumor-specific and-expressed lncRNA (GSVA-score), immune cell abundance (CIBERSORT scores of M1 macrophages, CD4 memory activated T cells, CD8 T cells and regulatory T cells), as well as clinical information (ECOG ≥ 1 (Y/N), liver metastasis (Y/N)).

We tested three different machine learning algorithms on our data to build a robust prediction model. Logistic regression with L1 regularization type was used as a baseline model. Another model that showed better performances than logistic regression and also provided positive and negative coefficients is the stochastic gradient descent. Finally, random forest was used for the final model. This method is very robust to outliers by averaging the scores of all the trees that comprehend the forest. Note that the feature importance that can be extracted from the random forest is based on the gini impurity decrease; hence, they are always positive as they are percentages.

To account for the limited sample size, we tested the model with 1000 different seeds for more robust and reliable scores. Furthermore, we integrated Bootstrap.632+ as an internal validation technique. This method provides information on how the model may behave in different datasets not involved in the training phase ^69^. Additionally, we repeatedly run the model, removing one of the five datasets that have RNA-Seq data respectively. The AUC did not change by more than 0.016 (2.14%) when UNC-108 was removed, and by 0.007 (0.94%) for the rest of the datasets, ensuring that our results are not driven by one of the datasets.

Finally, we tested the model in the independent validation cohort JAVELIN Bladder 100 trial. We were provided with processed gene expression and mutation call data. From the TPM matrices, we obtained the gene expression levels of selected genes (PD-1, PD-L1, CCND1), immune signatures using GSVA (IFN-γ, stroma/EMT, inflamed T-cells, TGF-β, and antigen-presenting machinery pathways), and immune cell abundances from CIBERSORT (M1 macrophages, CD4 memory activated T cells, CD8 T cells, and regulatory T cells). From the list of called mutations per patient, we obtain an estimate of TMB. Due to the lack of raw WES data and incomplete clinical information, we could not determine the number of non-stop mutations, the APOBEC-enrichment score, the status of liver metastasis, or the ECOG score of the patients. Moreover, raw RNA-Seq data would have been needed to compute the signature of tumor-specific lncRNAs. We used the TMB + RNA model (Figure 6A), the lncRNA signature was given a 0 value.

### Statistical analysis

Plots were mainly generated using R (version 4.1.2), Rstudio (version 1.4) and Python (version 3.8.6). When comparing continuous variables between the two response groups, significance levels were obtained using the Wilcoxon signed-rank test. We used the Bonferroni correction post-hoc to assess the significance of the effect of expression of single genes or gene signatures. As 15 gene signatures were tested, the adjusted alpha for gene signatures was 0.00033. For genes, the alpha for adjusted p-values it was 0.00029. When comparing ratios the Fisher exact test was employed. Pearson correlation was used to compare numeric variables.

## Data Availability

Suppl. file 1 contains Suppl. Figures and Tables indicated in the text. Suppl. file 2 contains a list of the variables investigated, descriptive statistics of the distribution of the variable values in different groups of patients, and numerical values related to the plots in the Figures. The code for the model and decision trees is available from the github repository https://github.com/EvolutionaryGenomics-GRIB/ML_Pipeline. The trained model can be executed at https://github.com/EvolutionaryGenomics-GRIB/BLCA_ICI_Response_Predictor.

## ACKNOWLEDGEMENTS

The work was supported by the following grants and agencies: 1. Research project PID2019-105595GB and PID2021-122726NB-I00 funded by MCIN/AEI/10.13039/501100011033 and by “ERDF: A way of making Europe”, by the “European Union”; 2. FIS PI22/00171, funded by Instituto de Salud Carlos III (ISCIII) and co-funded by the European Union and FEDER; 3. 2021SGR00042 by Generalitat de Catalunya; 4. Ayudas Fundación BBVA a Proyectos de Investigación Científica en Biomedicina 2021. LMB is funded by an INPhINIT PhD fellowship from “la Caixa” Foundation (ID 100010434), under the agreement LCF/BQ/DI21/11860060.

## AUTHOR CONTRIBUTIONS

L.M.B. and S.V. contributed equally to this work. J.P.B., M.M.A., S.V. and L.M.B. gathered and integrated the datasets of this study. M.E., S.V. and L.M.B. performed the computational analysis of RNA-seq data. L.M.B. conducted the computational analysis of WES data. S.V. and R.C. developed the machine learning models. M.M.A., J.P.B and J.B. contributed to the conceptualization of the study and design of experiments. M.M.A., L.M.B., S.V., and M.E. and wrote the manuscript with contributions from all co-authors.

## COMPETING INTERESTS

Potential conflicts of interest: J. Bellmunt has served in consulting or advisory roles for Astellas Pharma, AstraZeneca/MedImmune, Bristol Myers Squibb, Genentech, Novartis, Pfizer, Pierre Fabre, and the healthcare business of Merck KGaA, Darmstadt, Germany; has received travel and accommodation expenses from Ipsen, Merck & Co., Kenilworth, NJ, and Pfizer; reports patents, royalties, other intellectual property from UpToDate; reports stock and other ownership interests in Rainier Therapeutics; has received honoraria from UpToDate; and has received institutional research funding from Millennium, Pfizer, Sanofi, and the healthcare business of Merck KGaA, Darmstadt, Germany.

## Notes

### Summary of Updates

Results, Discussion and Methods section updated to include external model validation using the JAVELIN Bladder 100 trial cohort; Results section updated to contain correlation coefficients; Figure 6 revised; Supplemental files updated.

## REFERENCES

1. Bellmunt, J. et al. Pembrolizumab as Second-Line Therapy for Advanced Urothelial Carcinoma. N. Engl. J. Med. 376, 1015–1026 (2017).

2. Powles, T. et al. Avelumab Maintenance Therapy for Advanced or Metastatic Urothelial Carcinoma. N. Engl. J. Med. 383, 1218–1230 (2020).

3. Miao, D. et al. Genomic correlates of response to immune checkpoint blockade in microsatellite-stable solid tumors. Nat. Genet. 50, 1271–1281 (2018).

4. Nadal, R., Valderrama, B. P. & Bellmunt, J. Progress in systemic therapy for advanced-stage urothelial carcinoma. Nat. Rev. Clin. Oncol. 21, 8–27 (2024).

5. Snyder, A. et al. Contribution of systemic and somatic factors to clinical response and resistance to PD-L1 blockade in urothelial cancer: An exploratory multi-omic analysis. PLoS Med. 14, e1002309 (2017).

6. McGrail, D. J. et al. High tumor mutation burden fails to predict immune checkpoint blockade response across all cancer types. Ann. Oncol. 32, 661–672 (2021).

7. Davis, E. J. et al. Clinical Correlates of Response to Anti-PD-1–based Therapy in Patients With Metastatic Melanoma. J. Immunother. 42, 221–227 (2019).

8. Galsky, M. D. et al. Perioperative pembrolizumab therapy in muscle-invasive bladder cancer: Phase III KEYNOTE-866 and KEYNOTE-905/EV-303. *Futur*. Oncol. 17, 3137–3150 (2021).

9. Mariathasan, S. et al. TGFβ attenuates tumour response to PD-L1 blockade by contributing to exclusion of T cells. Nature 554, 544–548 (2018).

10. Robertson, A. G. et al. Comprehensive Molecular Characterization of Muscle-Invasive Bladder Cancer. Cell 171, 540–556.e25 (2017).

11. Kamoun, A. et al. A Consensus Molecular Classification of Muscle-invasive Bladder Cancer. Eur. Urol. 77, 420–433 (2020).

12. Comprehensive molecular characterization of urothelial bladder carcinoma. Nature 507, 315–322 (2014).

13. Rosenberg, J. E. et al. Atezolizumab in patients with locally advanced and metastatic urothelial carcinoma who have progressed following treatment with platinum-based chemotherapy: a single-arm, multicentre, phase 2 trial. Lancet 387, 1909–1920 (2016).

14. Kim, J. et al. The Cancer Genome Atlas Expression Subtypes Stratify Response to Checkpoint Inhibition in Advanced Urothelial Cancer and Identify a Subset of Patients with High Survival Probability. Eur. Urol. 75, 961–964 (2019).

15. Litchfield, K. et al. Meta-analysis of tumor-and T cell-intrinsic mechanisms of sensitization to checkpoint inhibition. Cell 184, 596–614.e14 (2021).

16. Damrauer, J. S. et al. Collaborative study from the Bladder Cancer Advocacy Network for the genomic analysis of metastatic urothelial cancer. Nat. Commun. 13, 6658 (2022).

17. Long, J. et al. A mutation-based gene set predicts survival benefit after immunotherapy across multiple cancers and reveals the immune response landscape. Genome Med. 14, 20 (2022).

18. Boll, L. M. et al. The impact of mutational clonality in predicting the response to immune checkpoint inhibitors in advanced urothelial cancer. Sci. Rep. 13, 15287 (2023).

19. Rose, T. L. et al. Fibroblast growth factor receptor 3 alterations and response to immune checkpoint inhibition in metastatic urothelial cancer: a real world experience. Br. J. Cancer 125, 1251–1260 (2021).

20. Martínez-Jiménez, F. et al. A compendium of mutational cancer driver genes. Nat. Rev. Cancer 20, 555–572 (2020).

21. Yang, Y. et al. Targeting ARHGEF12 promotes neuroblastoma differentiation, MYCN degradation, and reduces tumorigenicity. Cell. Oncol. 46, 133–143 (2023).

22. Galsky, M. D. et al. Impact of zumor mutation burden on nivolumab efficacy in second-line urothelial carcinoma patients: Exploratory analysis of the phase ii checkmate 275 study. Ann. Oncol. 28, v296–v297 (2017).

23. Reynisson, B., Alvarez, B., Paul, S., Peters, B. & Nielsen, M. NetMHCpan-4.1 and NetMHCIIpan-4.0: improved predictions of MHC antigen presentation by concurrent motif deconvolution and integration of MS MHC eluted ligand data. Nucleic Acids Res. 48, W449–W454 (2020).

24. Van Allen, E. M. et al. Genomic correlates of response to CTLA-4 blockade in metastatic melanoma. Science 350, 207–211 (2015).

25. Nassar, A. H. et al. A model combining clinical and genomic factors to predict response to PD-1/PD-L1 blockade in advanced urothelial carcinoma. Br. J. Cancer 122, 555–563 (2020).

26. Adib, E., et al. *CDKN2A* Alterations and Response to Immunotherapy in Solid Tumors. Clin. Cancer Res. 27, 4025–4035 (2021).

27. Steen, C. B., Liu, C. L., Alizadeh, A. A. & Newman, A. M. Profiling Cell Type Abundance and Expression in Bulk Tissues with CIBERSORTx. in 135–157 (Humana, New York, NY, 2020). doi:10.1007/978-1-0716-0301-7_7

28. Philips, G. K. & Atkins, M. Therapeutic uses of anti-PD-1 and anti-PD-L1 antibodies. Int. Immunol. 27, 39–46 (2015).

29. Büttner, R. et al. Programmed Death-Ligand 1 Immunohistochemistry Testing: A Review of Analytical Assays and Clinical Implementation in Non–Small-Cell Lung Cancer. J. Clin. Oncol. 35, 3867–3876 (2017).

30. Baxi, V. et al. Association of artificial intelligence-powered and manual quantification of programmed death-ligand 1 (PD-L1) expression with outcomes in patients treated with nivolumab ± ipilimumab. Mod. Pathol. 35, 1529–1539 (2022).

31. Shen, X. & Zhao, B. Efficacy of PD-1 or PD-L1 inhibitors and PD-L1 expression status in cancer: meta-analysis. BMJ 362, k3529 (2018).

32. McDermott, D. F. et al. Clinical activity and molecular correlates of response to atezolizumab alone or in combination with bevacizumab versus sunitinib in renal cell carcinoma. Nat. Med. 24, 749–757 (2018).

33. Fehrenbacher, L. et al. Atezolizumab versus docetaxel for patients with previously treated non-small-cell lung cancer (POPLAR): a multicentre, open-label, phase 2 randomised controlled trial. *Lancet (London*, England) 387, 1837–46 (2016).

34. Bindea, G. et al. Spatiotemporal Dynamics of Intratumoral Immune Cells Reveal the Immune Landscape in Human Cancer. Immunity 39, 782–795 (2013).

35. Ayers, M. et al. IFN-γ-related mRNA profile predicts clinical response to PD-1 blockade. J. Clin. Invest. 127, 2930–2940 (2017).

36. Cristescu, R. et al. Pan-tumor genomic biomarkers for PD-1 checkpoint blockade-based immunotherapy. Science 362, (2018).

37. Hernando-Calvo, A. et al. A pan-cancer clinical platform to predict immunotherapy outcomes and prioritize immuno-oncology combinations in early-phase trials. Med 4, 710–727.e5 (2023).

38. Benguigui, M. et al. Interferon-stimulated neutrophils as a predictor of immunotherapy response. Cancer Cell 42, 253–265.e12 (2024).

39. Thompson, J. C. et al. Gene signature of antigen processing and presentation machinery predicts response to checkpoint blockade in non-small cell lung cancer (NSCLC) and melanoma. J. Immunother. cancer 8, e000974 (2020).

40. Ouspenskaia, T. et al. Unannotated proteins expand the MHC-I-restricted immunopeptidome in cancer. Nat. Biotechnol. (2021). doi:10.1038/s41587-021-01021-3

41. Ruiz Cuevas, M.V., et al. Most non-canonical proteins uniquely populate the proteome or immunopeptidome. Cell Rep. 34, 108815 (2021).

42. Laumont, C. M. et al. Noncoding regions are the main source of targetable tumor-specific antigens. Sci. Transl. Med. 10, (2018).

43. Wang, L. et al. EMT-and stroma-related gene expression and resistance to PD-1 blockade in urothelial cancer. Nat. Commun. 9, 3503 (2018).

44. Sha, D. et al. Tumor Mutational Burden as a Predictive Biomarker in Solid Tumors. Cancer Discov. 10, 1808–1825 (2020).

45. Powles, T. et al. Avelumab maintenance in advanced urothelial carcinoma: biomarker analysis of the phase 3 JAVELIN Bladder 100 trial. Nat. Med. 27, 2200–2211 (2021).

46. Togashi, Y., Shitara, K. & Nishikawa, H. Regulatory T cells in cancer immunosuppression — implications for anticancer therapy. Nat. Rev. Clin. Oncol. 16, 356–371 (2019).

47. Boichard, A. et al. APOBEC-related mutagenesis and neo-peptide hydrophobicity: implications for response to immunotherapy. Oncoimmunology 8, 1550341 (2019).

48. Bellmunt, J. et al. Genomic Predictors of Good Outcome, Recurrence, or Progression in High-Grade T1 Non–Muscle-Invasive Bladder Cancer.Cancer Res. 80, 4476–4486 (2020).

49. Turajlic, S. et al. Insertion-and-deletion-derived tumour-specific neoantigens and the immunogenic phenotype: a pan-cancer analysis. Lancet. Oncol. 18, 1009–1021 (2017).

50. Zhang, J. et al. Cyclin D–CDK4 kinase destabilizes PD-L1 via cullin 3–SPOP to control cancer immune surveillance. Nature 553, 91–95 (2018).

51. Barczak, W. et al. Long non-coding RNA-derived peptides are immunogenic and drive a potent anti-tumour response. Nat. Commun. 14, 1078 (2023).

52. Xiong, D. et al. Immunogenomic Landscape Contributes to Hyperprogressive Disease after Anti-PD-1 Immunotherapy for Cancer. iScience 9, 258–277 (2018).

53. Chowell, D. et al. Improved prediction of immune checkpoint blockade efficacy across multiple cancer types. Nat. Biotechnol. 40, 499–506 (2022).

54. Liu, D. et al. Integrative molecular and clinical modeling of clinical outcomes to PD1 blockade in patients with metastatic melanoma. Nat. Med. 25, 1916–1927 (2019).

55. Griffin, J. et al. Verification of molecular subtyping of bladder cancer in the GUSTO clinical trial. J. Pathol. Clin. Res. 10, e12363 (2024).

56. Balar, A. V. & Weber, J. S. PD-1 and PD-L1 antibodies in cancer: current status and future directions. Cancer Immunol. Immunother. 66, 551–564 (2017).

57. Mayakonda, A., Lin, D.-C., Assenov, Y., Plass, C. & Koeffler, H. P. Maftools: efficient and comprehensive analysis of somatic variants in cancer. Genome Res. 28, 1747–1756 (2018).

58. Zack, T. I. et al. Pan-cancer patterns of somatic copy number alteration. Nat. Genet. 45, 1134–1140 (2013).

59. Rosenthal, R., McGranahan, N., Herrero, J., Taylor, B. S. & Swanton, C. deconstructSigs: delineating mutational processes in single tumors distinguishes DNA repair deficiencies and patterns of carcinoma evolution. Genome Biol. 17, 31 (2016).

60. Ewels, P. A. et al. The nf-core framework for community-curated bioinformatics pipelines. Nat. Biotechnol. 38, 276–278 (2020).

61. Szolek, A. et al. OptiType: precision HLA typing from next-generation sequencing data. Bioinformatics 30, 3310–6 (2014).

62. Jurtz, V. et al. NetMHCpan-4.0: Improved Peptide-MHC Class I Interaction Predictions Integrating Eluted Ligand and Peptide Binding Affinity Data. J. Immunol. 199, 3360–3368 (2017).

63. Dobin, A. et al. STAR: ultrafast universal RNA-seq aligner. Bioinformatics 29, 15–21 (2013).

64. Wang, L., Wang, S. & Li, W. RSeQC: quality control of RNA-seq experiments. Bioinformatics 28, 2184–2185 (2012).

65. Liao, Y., Smyth, G. K. & Shi, W. featureCounts: an efficient general purpose program for assigning sequence reads to genomic features. Bioinformatics 30, 923–30 (2014).

66. Hänzelmann, S., Castelo, R. & Guinney, J. GSVA: gene set variation analysis for microarray and RNA-seq data. BMC Bioinformatics 14, 7 (2013).

67. Tokunaga, R. et al. 12-Chemokine signature, a predictor of tumor recurrence in colorectal cancer. Int. J. Cancer 147, 532–541 (2020).

68. Chong, C. et al. Integrated proteogenomic deep sequencing and analytics accurately identify non-canonical peptides in tumor immunopeptidomes. Nat. Commun. 11, 1293 (2020).

69. Efron, B. & Tibshirani, R. Improvements on Cross-Validation: The 632+ Bootstrap Method. J. Am. Stat. Assoc. 92, 548–560 (1997).

