## Supplementary File 1 for "Predicting immunotherapy response in advanced bladder cancer: a meta-analysis of six independent cohorts"

| | NR *N=303* | R *N=163* | p.overall |
|---|---|---|---|
| ***RECIST:*** | | | ***<0.001*** |
| Complete Response | - | 56 (34.4%) | |
| Partial Response | - | 107 (65.6%) | |
| Progressive Disease | 303 (100%) | - | |
| ***Sex:*** | | | ***0.196*** |
| female | 73 (24.1%) | 30 (18.4%) | |
| male | 230 (75.9%) | 133 (81.6%) | |
| **ECOG** | **0.80 (0.72)** | **0.66 (0.77)** | **0.060** |
| **ECOG ≥ 1:** | | | **0.006** |
| No | 186 (66.0%) | 75 (51.7%) | |
| Yes | 96 (34.0%) | 70 (48.3%) | |
| ***Age*** | ***65.6 (10.6)*** | ***67.1 (10.3)*** | ***0.319*** |
| ***Smoker:*** | | | ***0.143*** |
| No | 86 (30.4%) | 31 (23.0%) | |
| Yes | 197 (69.6%) | 104 (77.0%) | |
| ***Alive:*** | | | ***<0.001*** |
| No | 92 (76.0%) | 23 (29.5%) | |
| Yes | 29 (24.0%) | 55 (70.5%) | |
| ***OS*** | ***119 (384)*** | ***269 (455)*** | ***<0.001*** |
| ***Liver metastasis:*** | | | ***0.001*** |
| No | 165 (66.8%) | 79 (85.9%) | |
| Yes | 82 (33.2%) | 13 (14.1%) | |
| ***ICI Drug:*** | | | ***0.043*** |
| anti-CTLA-4 + anti-PD-1/PD-L1 | 0 (0.00%) | 1 (0.83%) | |
| anti-PD-1/anti-PD-L1 | 10 (3.79%) | 12 (9.92%) | |
| atezolizumab | 202 (76.5%) | 88 (72.7%) | |
| avelumab | 2 (0.76%) | 3 (2.48%) | |
| durvalumab | 4 (1.52%) | 1 (0.83%) | |
| nivolumab | 5 (1.89%) | 0 (0.00%) | |
| pembrolizumab | 41 (15.5%) | 16 (13.2%) | |

**Sup. Table 1. Clinical and demographic information by ICI response groups (N=466).** Table of all patients with response status complete or partial response (R) or non-response (NR). Table generated with compareGroups R package using the default tests (t-test and ANOVA for continuous normal-distributed, $X^2$-test for categorical variables, pairwise comparisons for more than 2 groups of continuous variables with with Pearson test and for categorical with Mantel-Haenszel test, in both cases applying the Turkey method to adjust for multiple testing.

| Mutation classification | Total (N=318) | R (N=89) | NR (N=144) |
|---|---|---|---|
| **Frame_Shift_Del** | 1373 | 673 | 34 |
| **Frame_Shift_Ins** | 507 | 197 | 178 |
| **In_Frame_Del** | 277 | 123 | 93 |
| **In_Frame_Ins** | 19 | 3 | 13 |
| **Missense_Mutation** | 55652 | 24423 | 18006 |
| **Nonsense_Mutation** | 5179 | 2225 | 1696 |
| **Nonstop_Mutation** | 85 | 40 | 28 |
| **Splice_Site** | 1102 | 428 | 414 |
| **Translation_Start_Site** | 81 | 35 | 27 |
| **Total** | **64275** | **28147** | **20849** |
| Mean number of mutations | 202 | 316 | 144 |

**Sup. Table 2. Abundance of different types of non-synonymous mutations.** WES data from the tumors of 318 patients was obtained using the same pipeline. The cohorts were IMvigor210, MIAO-2018, HdM-BLCA-1 and SNY-2017. R: responder; NR: non-responder.

| | | | | | | | | | | | |
|---|---|---|---|---|---|---|---|---|---|---|---|
| ***TTN*** | ***TP53*** | ***KMT2D*** | ***MUC16*** | ***ARID1A*** | ***MACF1*** | ***ELF3*** | ***KDM6A*** | ***FGFR3*** | ***SYNE1*** | ***CCDC168*** | ***DST*** |
| 254 | 139 | 114 | 95 | 86 | 71 | 67 | 59 | 58 | 57 | 56 | 54 |
| ***PLEC*** | ***FAT1*** | ***KMT2C*** | ***RNF213*** | ***FAT4*** | ***HMCN1*** | ***OBSCN*** | ***XIRP2*** | ***RB1*** | ***ABCA13*** | ***ERBB3*** | ***PIK3CA*** |
| 52 | 50 | 50 | 50 | 49 | 49 | 48 | 46 | 44 | 43 | 43 | 43 |
| ***ZFHX4*** | ***CSMD3*** | ***MUC4*** | ***PCLO*** | ***CHD9*** | ***RYR2*** | ***CREBBP*** | ***FSIP2*** | ***KMT2A*** | ***PKHD1L1*** | ***EP300*** | ***SPTAN1*** |
| 43 | 42 | 42 | 42 | 41 | 41 | 40 | 40 | 40 | 40 | 39 | 39 |
| ***ZFP36L1*** | ***ADGRV1*** | ***LRP1B*** | ***RYR3*** | ***USH2A*** | ***HERC1*** | ***ANK2*** | ***ATM*** | ***DNAH11*** | ***MKI67*** | ***AHNAK*** | ***DNAH5*** |
| 39 | 38 | 38 | 37 | 37 | 36 | 34 | 34 | 34 | 34 | 33 | 33 |
| ***KIAA1109*** | ***APOB*** | ***DNAH17*** | ***ERBB2*** | ***FBN2*** | ***MYH9*** | ***SPEN*** | ***SYNE2*** | ***UBR4*** | ***ANKRD11*** | ***MYCBP2*** | ***SPTA1*** |
| 33 | 32 | 32 | 32 | 32 | 32 | 32 | 32 | 32 | 31 | 31 | 31 |
| ***TSC1*** | ***CEP350*** | ***FLG*** | ***RYR1*** | ***AKAP9*** | ***BIRC6*** | ***LRP2*** | ***PRKDC*** | ***PTPN13*** | ***ANKRD17*** | ***CMYA5*** | ***CSMD1*** |
| 31 | 30 | 30 | 30 | 29 | 29 | 29 | 29 | 29 | 28 | 28 | 28 |
| ***DNAH6*** | ***DNAH8*** | ***PCNT*** | ***PKHD1*** | ***SACS*** | ***ALMS1*** | ***APC*** | ***ATR*** | ***CDKN1A*** | ***CEP192*** | ***DNAH9*** | ***FAM186A*** |
| 28 | 28 | 28 | 28 | 28 | 27 | 27 | 27 | 27 | 27 | 27 | 27 |
| ***FAT3*** | ***LAMA5*** | ***NIPBL*** | ***TPR*** | ***CDH23*** | ***DCHS2*** | ***DIDO1*** | ***DNAH10*** | ***DYNC2H1*** | ***KMT2B*** | ***NEB*** | ***NRXN1*** |
| 27 | 27 | 27 | 27 | 26 | 26 | 26 | 26 | 26 | 26 | 26 | 26 |
| ***SMARCA4*** | ***ANK3*** | ***COL6A3*** | ***LRRK2*** | ***MUC3A*** | ***NBEAL1*** | ***PCDH17*** | ***PDZD2*** | ***SRCAP*** | | | |
| 26 | 25 | 25 | 25 | 25 | 25 | 25 | 25 | 25 | | | |

**Sup. Table 3. Most frequently mutated genes with the number of non-synonymous mutations combined for the four WES cohorts (IMvigor210, MIAO-2018, HdM-BLCA-1 and SNY-2017).** Genes are sorted from most to fewest mutations found for the gene, only showing genes with more than 24 somatic mutations.

| Model | Split-sample AUC | Bootstrap .632+ AUC |
|---|---|---|
| Baseline model complete data | 0.678 | 0.711 |
| Baseline model tumor-infiltrated data | 0.704 | 0.737 |
| Baseline model not-tumor-infiltrated data | 0.652 | 0.664 |
| Complete model complete data | 0.761 | 0.789 |
| Complete model tumor-infiltrated data | 0.793 | 0.817 |
| Complete model not-tumor-infiltrated data | 0.639 | 0.646 |
| TMB + RNA model complete data | 0.747 | 0.773 |
| TMB + RNA model tumor-infiltrated data | 0.769 | 0.792 |
| TMB + RNA model not-tumor-infiltrated data | 0.647 | 0.643 |

**Sup. Table 4. AUC values for different random forest models.** Baseline model corresponds to using only the variable TMB. Complete model is obtained with 17 variables, including mutation, gene expression and clinical variables. TMB + RNA model includes TMB and gene expression variables. The area under the curve (AUC) is the average of 1000 runs. Data is shown for the split-sample method and the internal bootstrapping method.

| Luminal Papillary<br>N=71; AUC (train/test) = 0.6544 | Luminal Infiltrated<br>N=138; AUC(train/test) = 0.8022 | Basal squamous<br>N=91; AUC (train/test) = 0.8223 |
|---|---|---|
| Macrophages M1: 0.109<br>TMB_zscore: 0.106<br>PD1.zscore: 0.093<br>IFNg_Ayers.GSVA: 0.074<br>T cells CD8: 0.069<br>CCND1: 0.066<br>Stroma_EMT.GSVA: 0.063<br>PDL1.zscore: 0.059<br>APM_8.GSVA: 0.058<br>TGF_beta.GSVA: 0.049<br>T cells CD4 memory activated: 0.048<br>HLA-I.GSVA: 0.047<br>T_cell_inflamed.GSVA: 0.044<br>T cells regulatory (Tregs): 0.042<br>t.spec.lncRNA.GSVA: 0.038<br>ECOG_1_or_larger: 0.017<br>Liver.Metastasis_Y: 0.017 | TMB_zscore: 0.169<br>Macrophages M1: 0.13<br>IFNg_Ayers.GSVA: 0.086<br>T_cell_inflamed.GSVA: 0.066<br>t.spec.lncRNA.GSVA: 0.063<br>TGF_beta.GSVA: 0.062<br>Stroma_EMT.GSVA: 0.06<br>APM_8.GSVA: 0.058<br>HLA-I.GSVA: 0.05<br>CCND1: 0.048<br>PDL1.zscore: 0.046<br>PD1.zscore: 0.044<br>T cells CD4 memory activated: 0.042<br>T cells CD8: 0.037<br>T cells regulatory (Tregs): 0.022<br>ECOG_1_or_larger: 0.009<br>Liver.Metastasis_Y: 0.007 | TMB_zscore: 0.146<br>Macrophages M1: 0.085<br>PD1.zscore: 0.084<br>IFNg_Ayers.GSVA: 0.083<br>HLA-I.GSVA: 0.08<br>CCND1: 0.075<br>APM_8.GSVA: 0.064<br>T cells CD8: 0.063<br>T cells regulatory (Tregs): 0.059<br>PDL1.zscore: 0.053<br>t.spec.lncRNA.GSVA: 0.041<br>Stroma_EMT.GSVA: 0.04<br>TGF_beta.GSVA: 0.04<br>T_cell_inflamed.GSVA: 0.036<br>T cells CD4 memory activated: 0.032<br>Liver.Metastasis_Y: 0.011<br>ECOG_1_or_larger: 0.008 |

**Sup. Table 5. Random forest models to predict response to ICI in different bladder cancer subtypes**. The variables selected were TMB z-score, clinical variables (ECOG, liver metastasis) and RNA-Seq-derived variables.

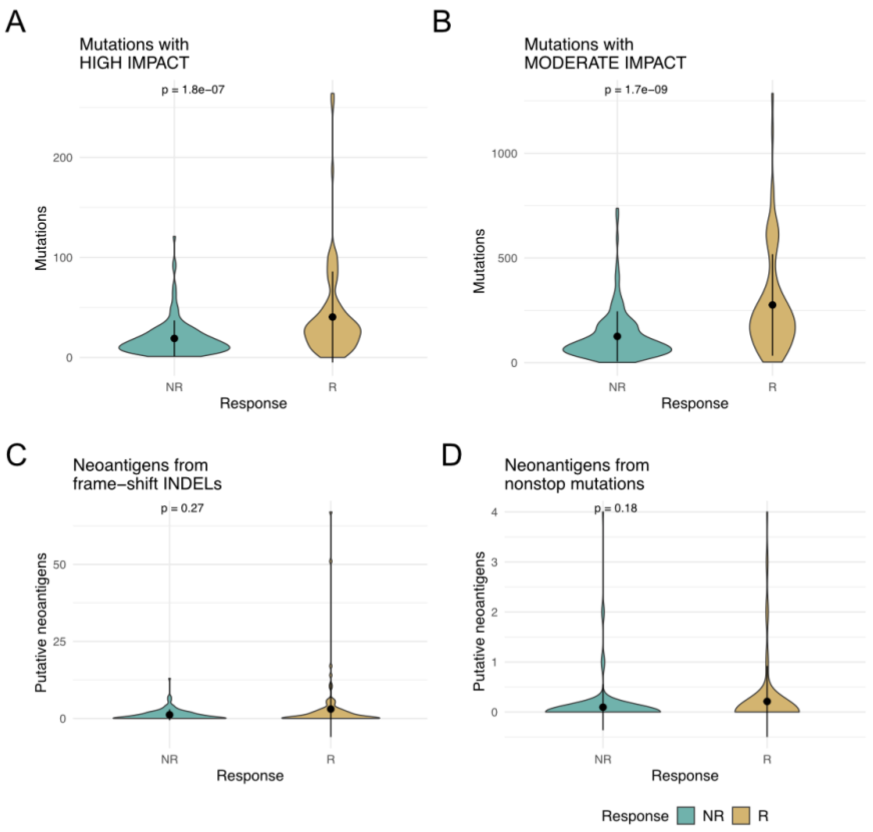


**Sup. Figure 1. Extended list of mutation-based variables. A.** Responders have higher number of mutations annotated as HIGH impact. **B.** The number of MODERATE impact mutations is also significantly higher in responders than in non-responders. **C.** The number of putative neoantigens resulting from frameshift insertions and deletions (fsINDELs) is not significantly different between responders and non-responders. **D.** The number of putative neoantigens resulting from nonstop mutations was not found to be significantly associated with response. N=236. P-values obtained by two-sample Wilcoxon test: R: responders, NR: non-responders. Mutation impact was obtained from ENSEMBL vep annotation (version 104). Putative binding peptides were predicted by applying a threshold of 500nM IC50 binding affinity in NetMHCpan 4.0.

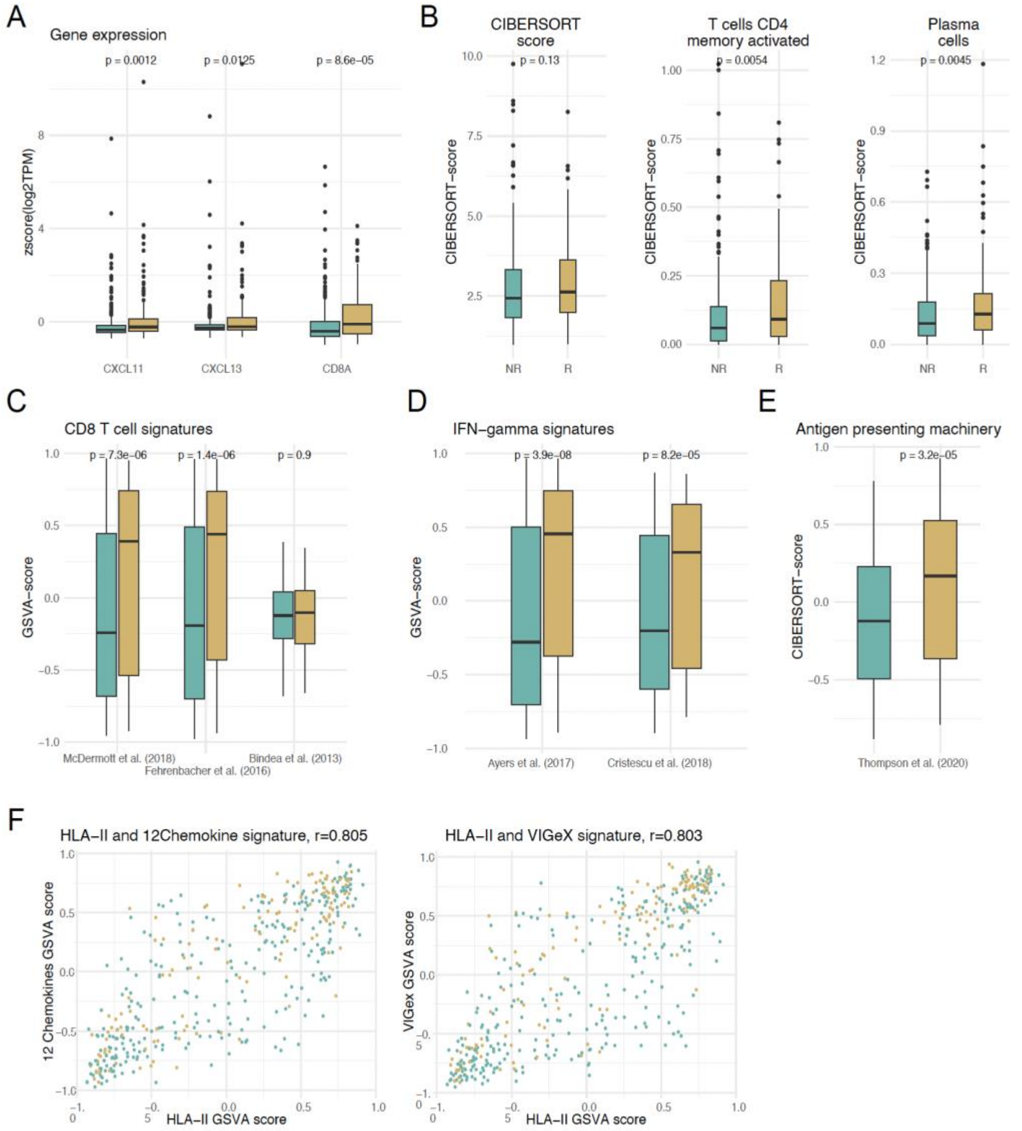


**Sup. Figure 2. Extended list of markers of immune activity in the tumor microenvironment. A.** Responders have higher gene expression of chemokines and CD8A compared to non-responders. **B.** Although responders do not show a higher general immune cell infiltration following the CIBERSORT absolute score, they have higher infiltration of cells related to an anti tumor response by the immune system. **C.** Responders are enriched in three out of four CD8 T cell signatures. **D.** Responders are enriched in IFN-gamma gene signatures. **E.** Responders are enriched in genes related to the antigen presenting machinery. **F.** High correlation between signatures of immune activation and HLA-II expression. N=420. P-values obtained by two-sample Wilcoxon test: R: responders, NR: non-responders. A full list of the included genes and sources for the gene signatures is provided in supplementary file 2.

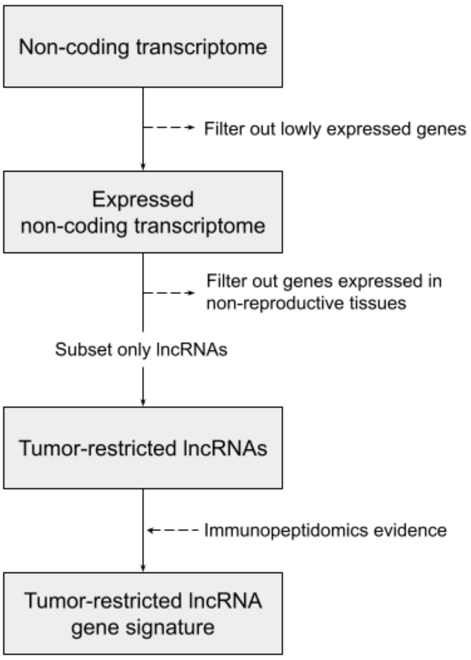


**Sup. Figure 3. Workflow overview of the identification of a tumor-restricted lncRNA gene signature.** Designed approach to detect tumor-restricted long non-coding RNAs and processed pseudogenes, collectively named lncRNAs. Cohorts were treated independently.

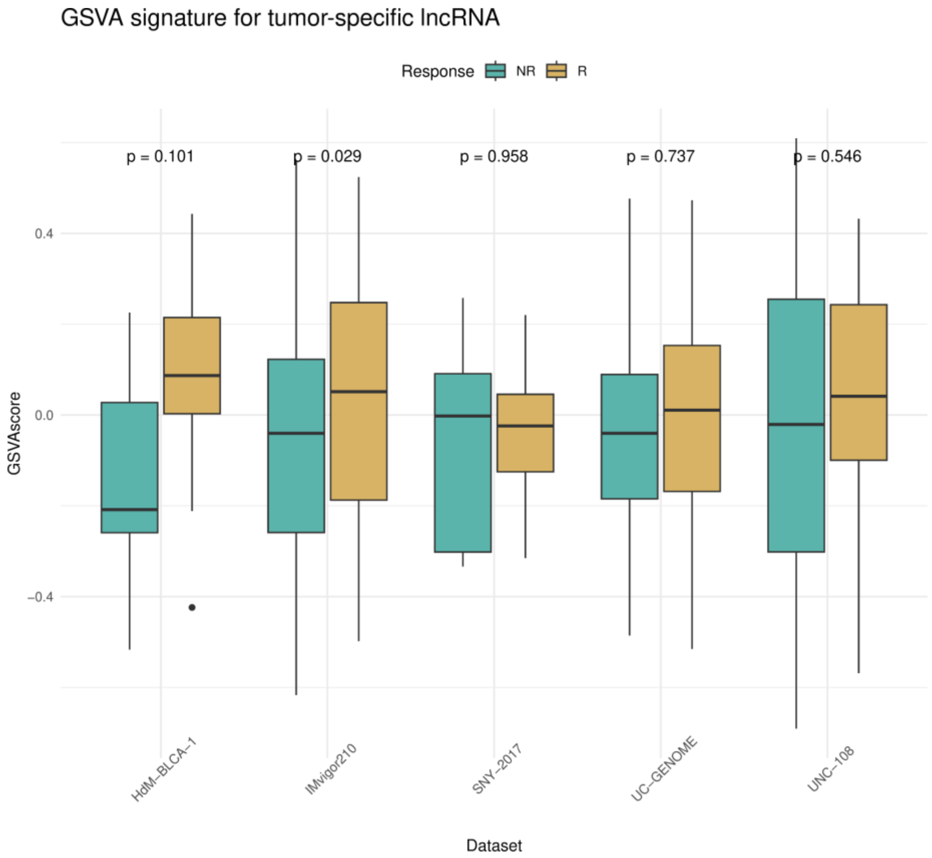


**Sup. Figure 4. Tumor-specific long non-coding RNAs gene signature.** GSVA signature score based on tumor-specific lncRNAs with immunopeptidomics evidence per dataset. P-values obtained by two-sample Wilcoxon test. R: responders, NR: non-responders.

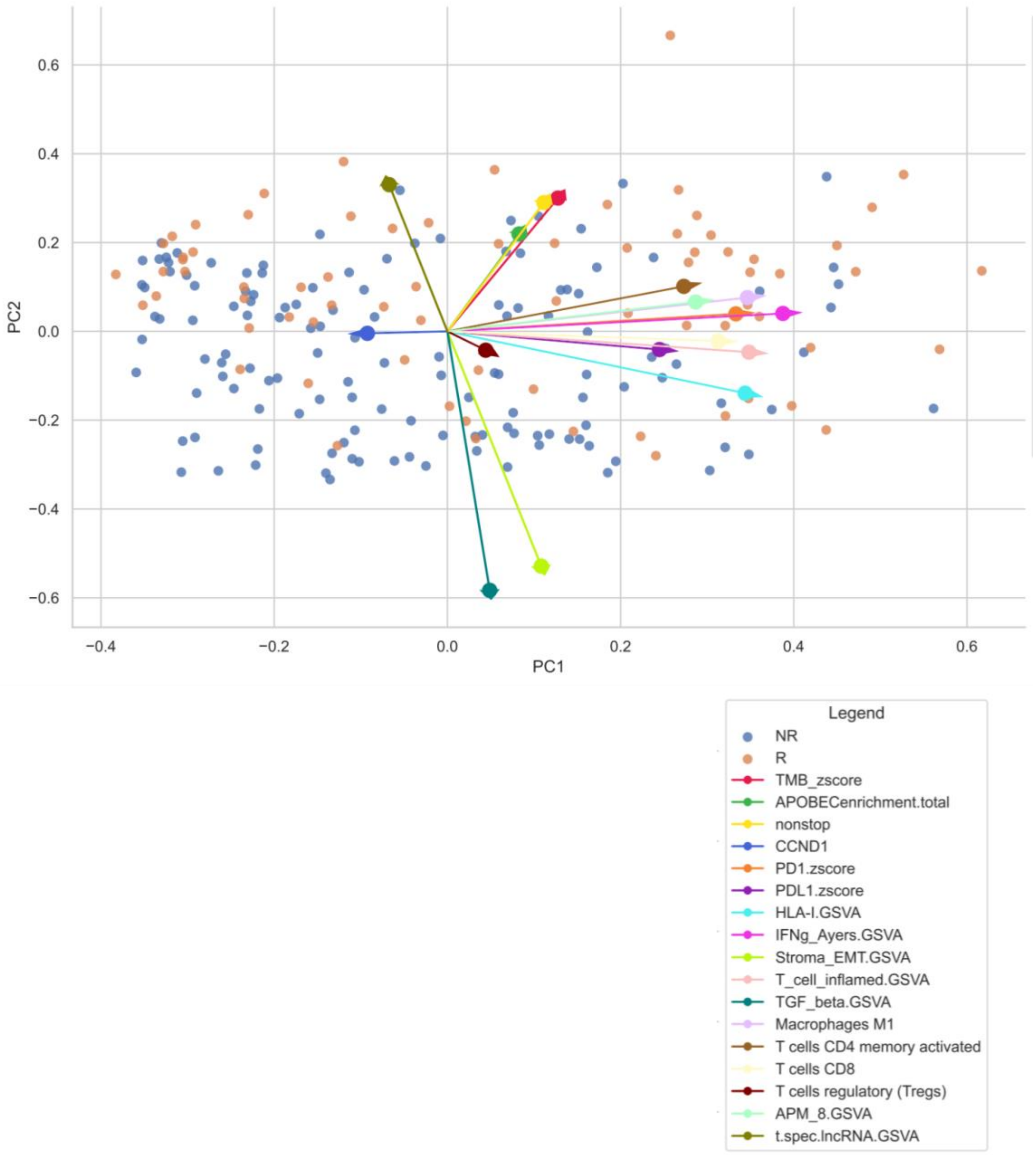


**Sup. Figure 5.** Principal component analysis of mutation and expression variables (N=205). Patients, colored by response, are represented by the smaller points in the plot, while variables are represented as vectors. The orientation and length of the vectors denote the relationship between variables and observations. Variables closer together in the biplot indicate higher correlation.

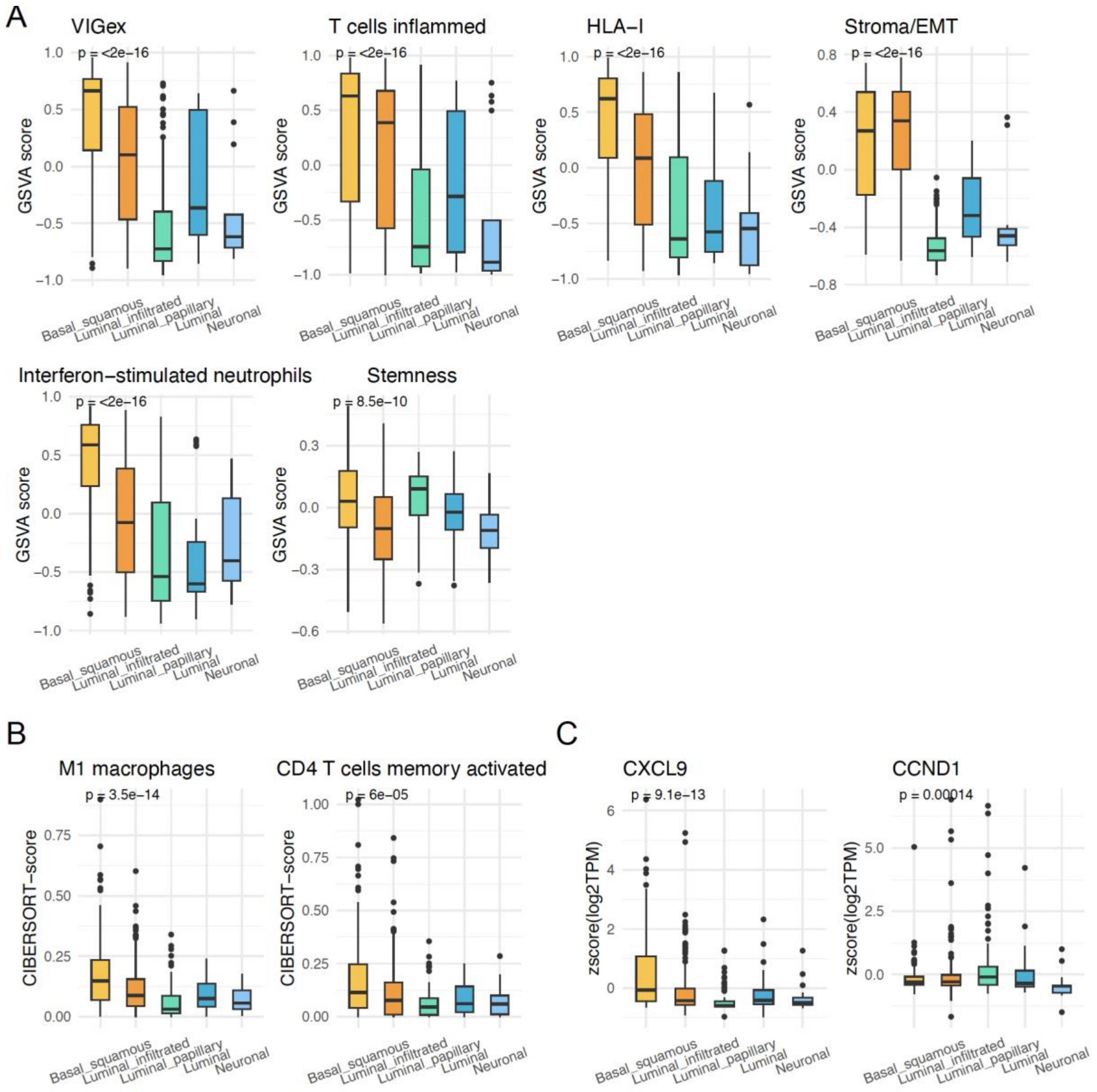


**Sup. Figure 6. Extended list of immune biomarkers in the five different TCGA subtypes. A.** Tumors of the basal-squamous subtype have the highest values of gene signatures related to immune activation as well as pathways of immune evasion such as Stroma/EMT. **B.** Basal-squamous is found to have the highest infiltration of CD8 T cells and M1 macrophages. **C.** Basal-squamous has the highest *CXCL9* expression. Luminal-papillary has the highest mean of *CCND1* expression. A full list of the included genes and sources for the gene signatures is provided in supplementary file 2.

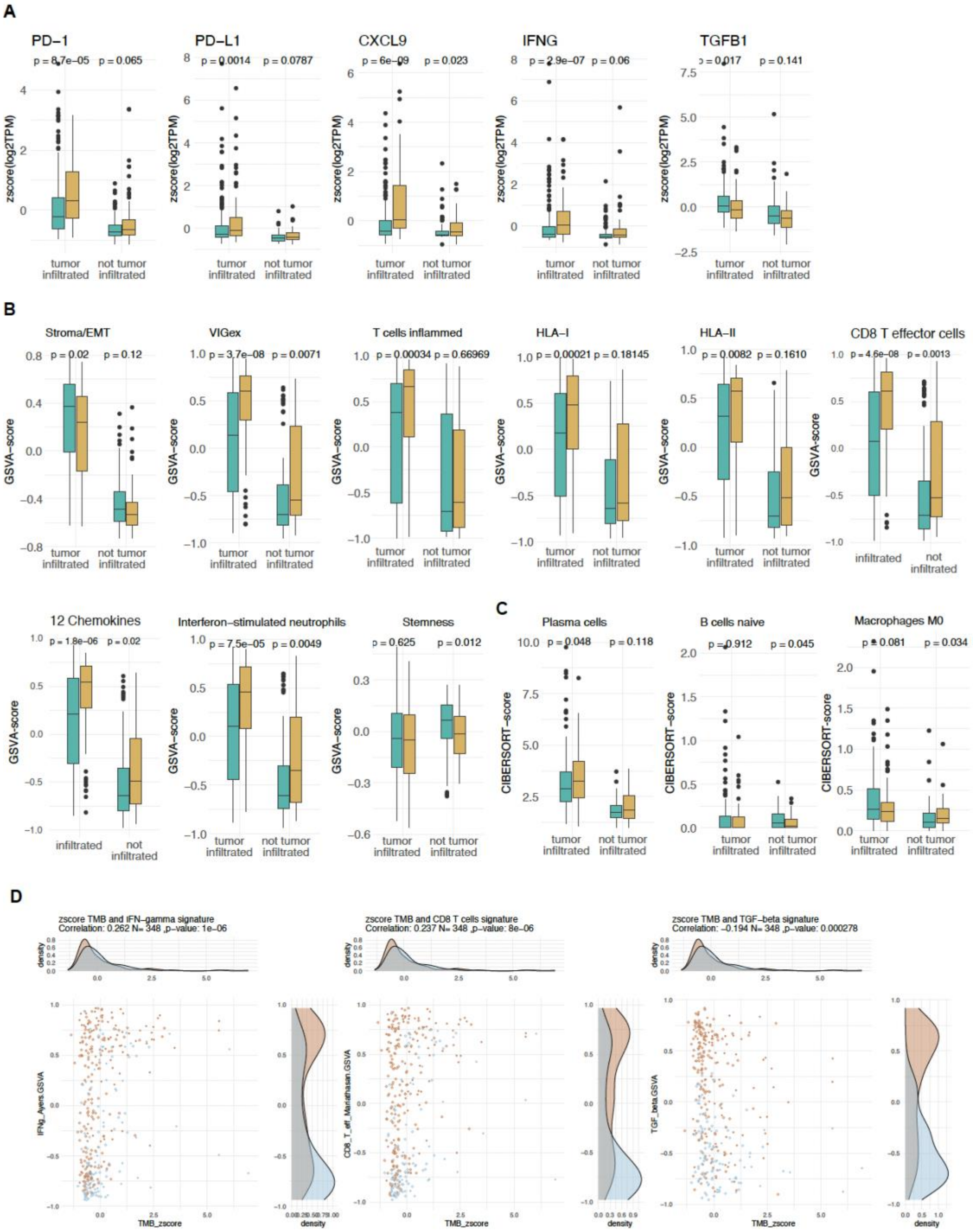


**Sup. Figure 7. Extended list of immune biomarkers in the immune-infiltrated and non-infiltrated subtypes. A.** Responders of the immune-infiltrated subtype have a higher mean expression of TGFB1 than non-responders. No difference was observed for the not-infiltrated subtype. **B**. Data for gene signatures. **C.** Data for immune cell deconvolution analysis. A full list of the included genes and sources for the gene signatures is provided in supplementary data 1.**D**. Relationship between TMB and immune-related variables, in immune-infiltrated subtypes (red) and non-immune-infiltrated subtypes (blue).

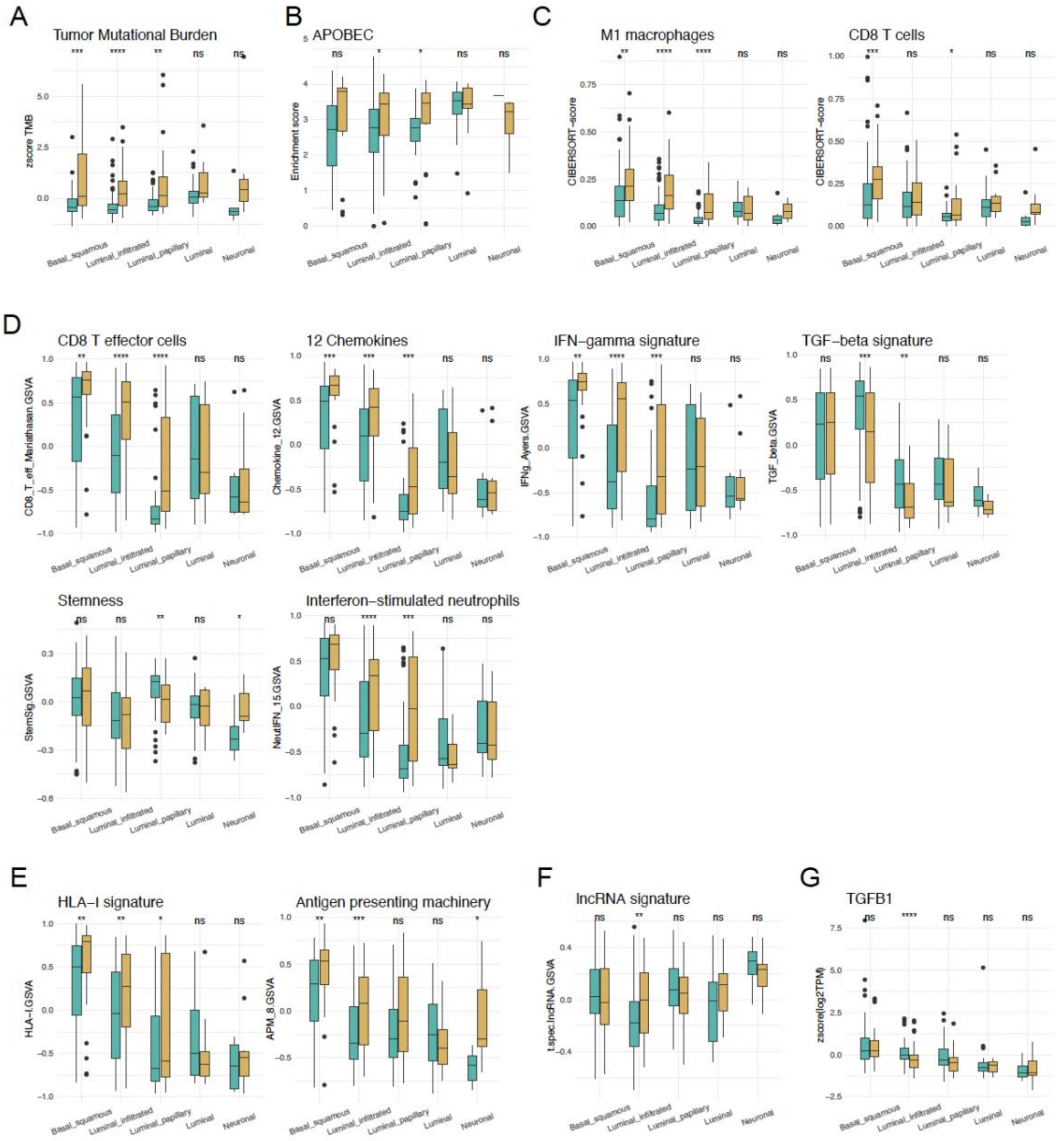


**Sup. Figure 8. Extended list of mutational and immune biomarkers in responders and non-responders by TCGA subtypes. A.** Relationship between TMB and subtype/response. **B.** Relationship between APOBEC enrichment and subtype/response. **C.** Relationship between immune cells and subtype/response. **D.** Relationship between immune activation and suppression signatures and subtype/response. **E.** Relationship between antigen presenting machinery (APM) pathway and subtype/response **F.** Relationship between the tumor specific lncRNA signature and subtype/response. **G.** Relationship between TGFB1 gene expression and subtype/response. N=420. P-values obtained by two-sample Wilcoxon test ns: p > 0.05, *: p <= 0.05, **: p <= 0.01, ***: p <= 0.001. R: responders, NR: non-responders. A full list of the included genes and sources for the gene signatures shown is provided in supplementary file 2.

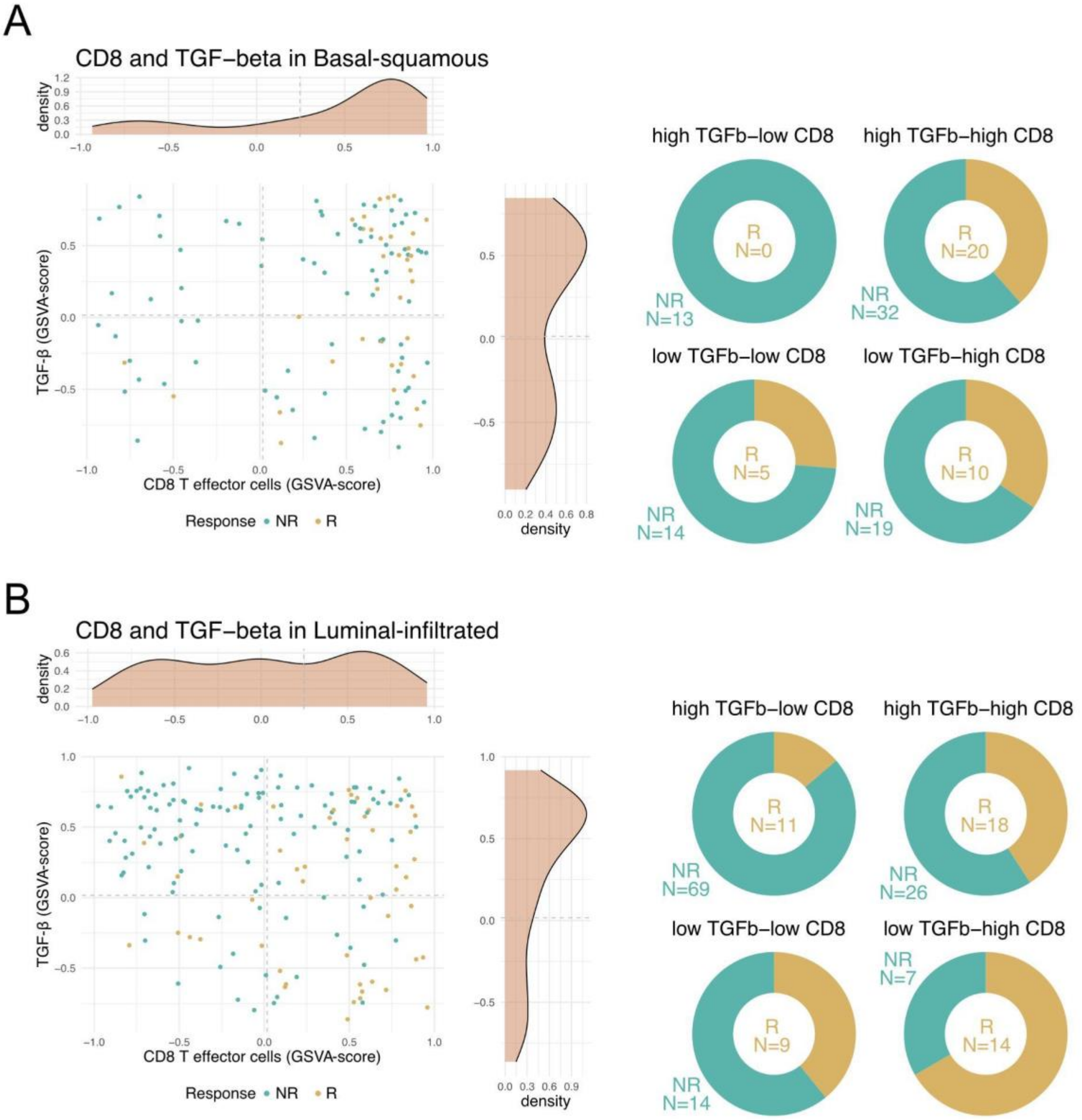


**Sup Figure 9. Relationship between CD8 T cell and TGF- gene expression in immune-infiltrated subtypes.** Relationship between CD8 T cell and TGF-β gene expression by the immune-infiltrated TCGA subtypes basal-squamous (**A**) and luminal-infiltrated (**B**). Immune-infiltrated samples tend to have high CD8+ T cell abundance, and in many cases also high levels of the TGF-β signature. The basal-squamous has 38.46% responders with high CD8+ T cell infiltration and high TGF-β compared to 34.48% with high CD8+ T cell infiltration but low TGF-β values (Fisher's Exact test p-value = 0.81). In the luminal-infiltrated there were 66.67% responders with high CD8+ T cell infiltration and high TGF-β compared to 40.91% responders with high CD8+ T cell infiltration but low TGF-β values (Fisher's Exact test p-value = 0.066). Dashed line marks the optimal cutoff for each gene signature obtained by ROC (TGF-β: 0.0163; CD8 t effector cells: 0.246).

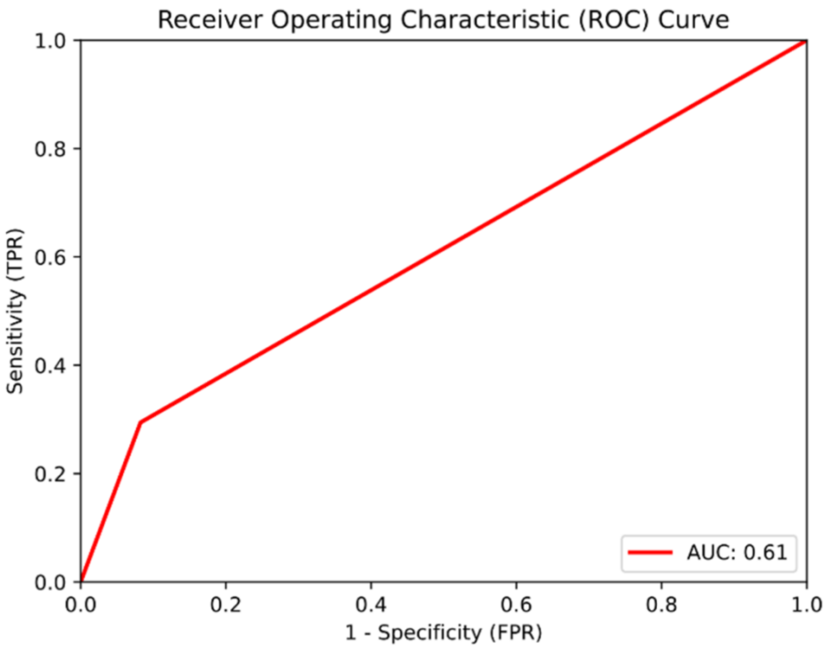


**Sup. Figure 10.** ROC curve obtained using a 10 mutations/Mb model. In this model tumors with less than 10 mutations/Mb are predicted to be non-responders.

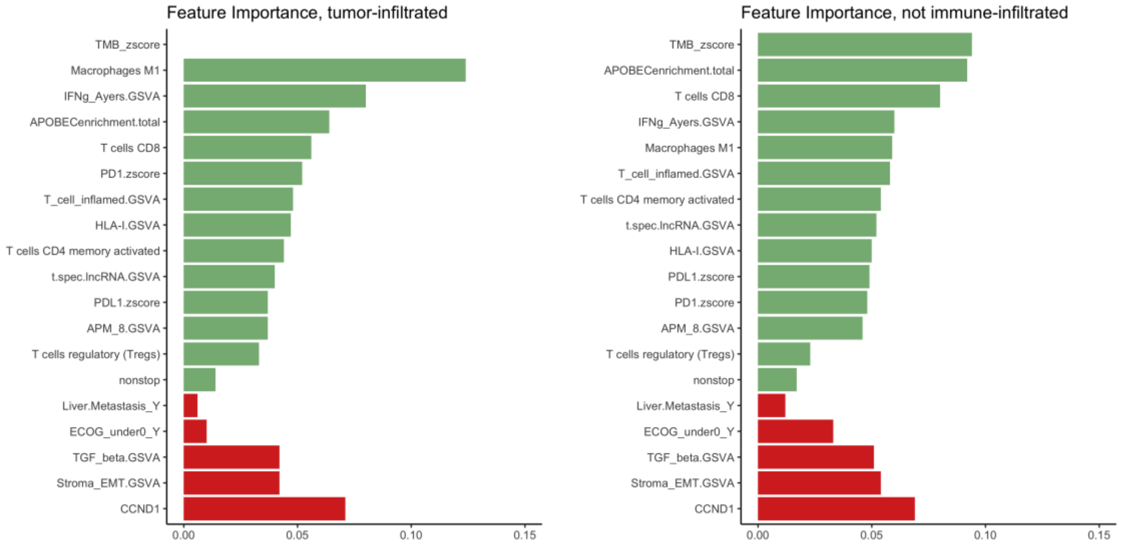


**Sup. Figure 11. Feature importance of all variables in the model.** The data is shown for immune-infiltrated subtypes (N=137) and not-immune infiltrated subtypes (N=68). Bars are colored according to their association with therapy response (green, higher in responders; red, higher in non-responders).

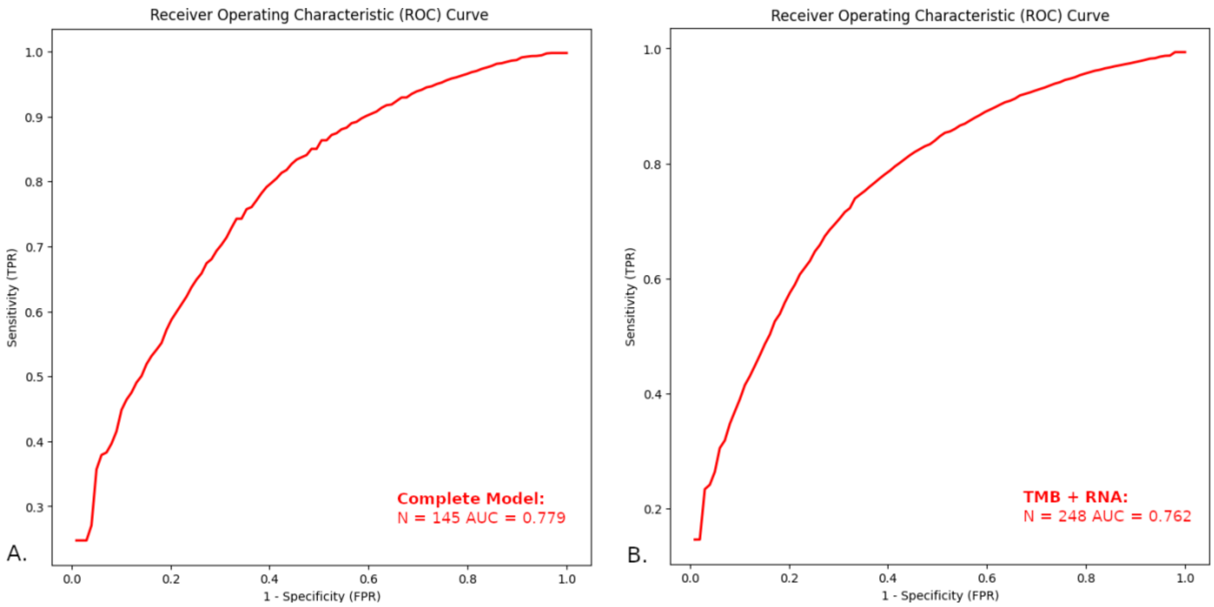


**Sup. Figure 12. Performance of the prediction model when undersampling.** The models were built with an equal number of responder and non-responders to assess performance without group imbalance. **(A)** Shows results for the Complete model and **(B)** reports the results for the TMB + RNA Model (TMB z-score and RNA-Seq-derived variables).

A.

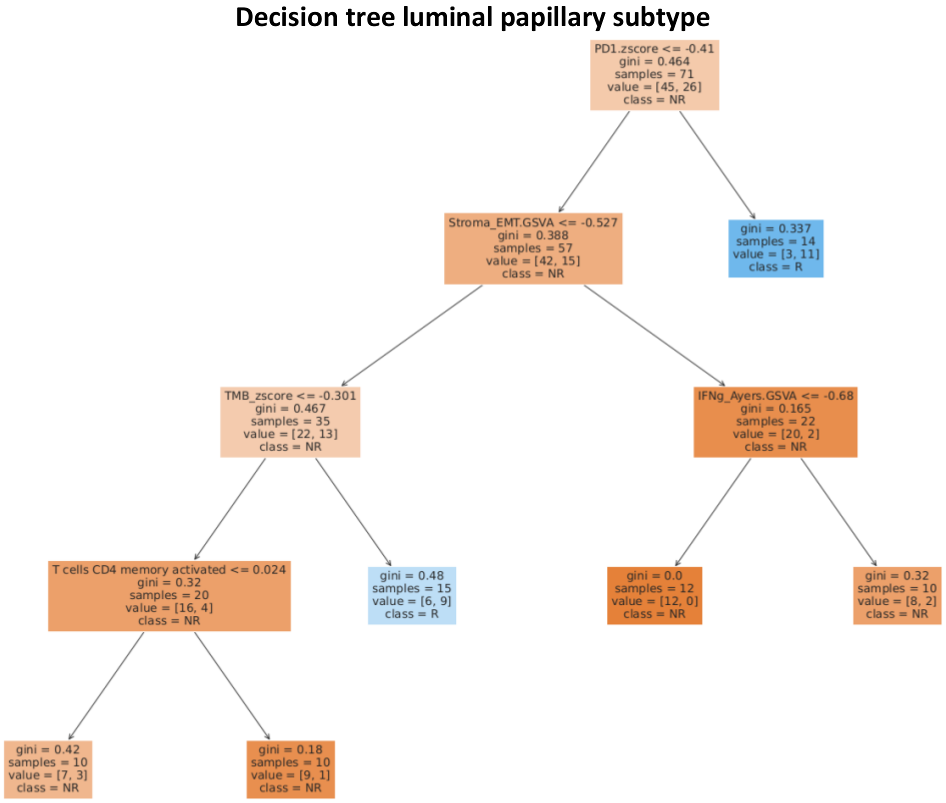


B.

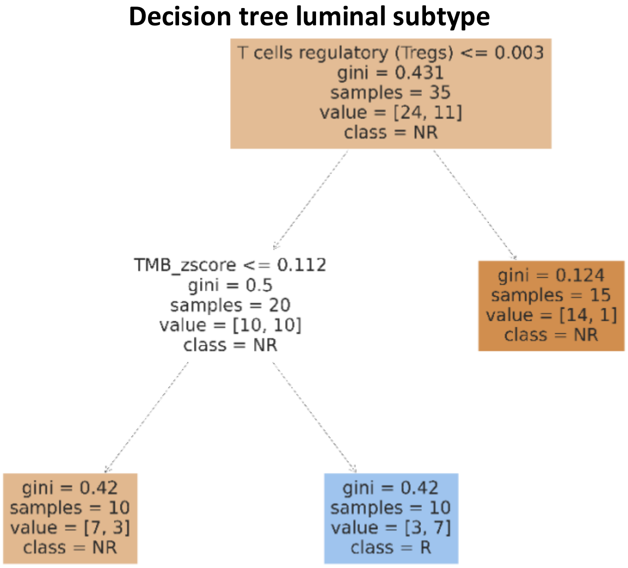


**Sup. Figure 13. Decision trees of non-immune-infiltrated subtypes.** Decision trees are created to better understand which features drive R/NR in our data stratifying by luminal-papillary (**A**) and luminal (**B**) TCGA Subtypes. The variables considered were TMB z-score and RNA-Seq-derived variables. The number of samples was 71 for luminal papillary and 35 for luminal. Number of leafs was set to 10.

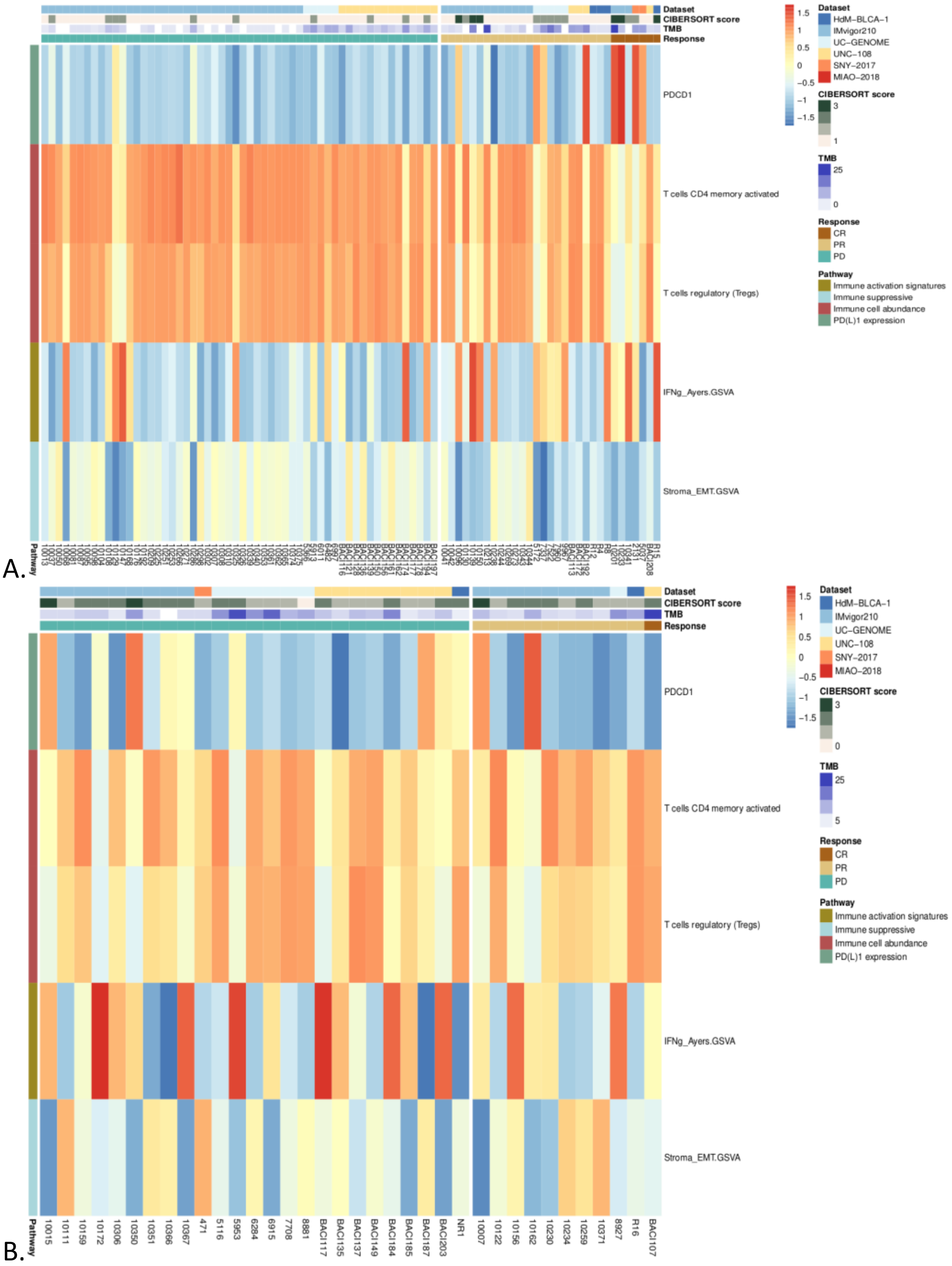


**Sup. Figure 14.** Heatmap of selected immune activation and suppression markers by non-immune infiltrated subtypes for luminal-papillary (A) and luminal (B). The five immune markers are taken from the decision trees in Sup. Figure 9.

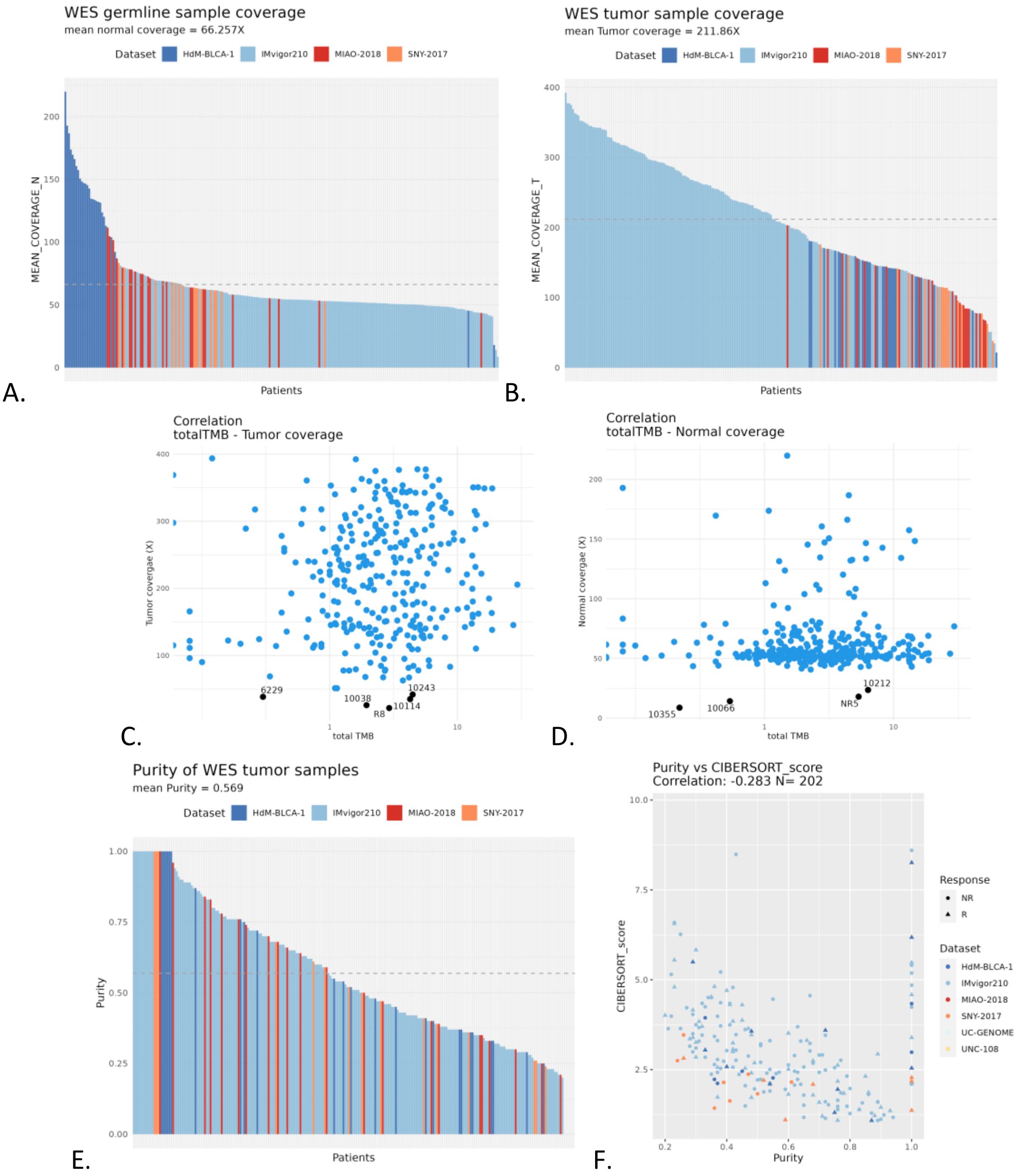


**Sup. Figure 15. Quality control of normal and tumor whole exome sequencing (WES) samples. A.** Coverage depth of germline WES samples. Samples from HdM-BLCA-1 were covered with a higher depth compared to the other three datasets. **B.** Coverage depth of WES from tumor samples. Samples of IMvigor210 have the highest coverage. **C.** Scatterplot showing the correlation between tumor mutational burden and tumor sample depth. Samples with a coverage > 20X are marked in black. As they don't fall into the 5% of samples with lowest mutation burden, they were not excluded in the analysis. **D.** Patients with a normal sample of low coverage don't show an unexpectedly high mutational burden and were therefore not excluded from the analysis. **E.** Purity estimates of tumor samples obtained from WES data using ASCAT. **F.** Correlation between WES tumor sample purity and deconvolution value obtained from CIBERSORT using tumor RNA-Seq data.

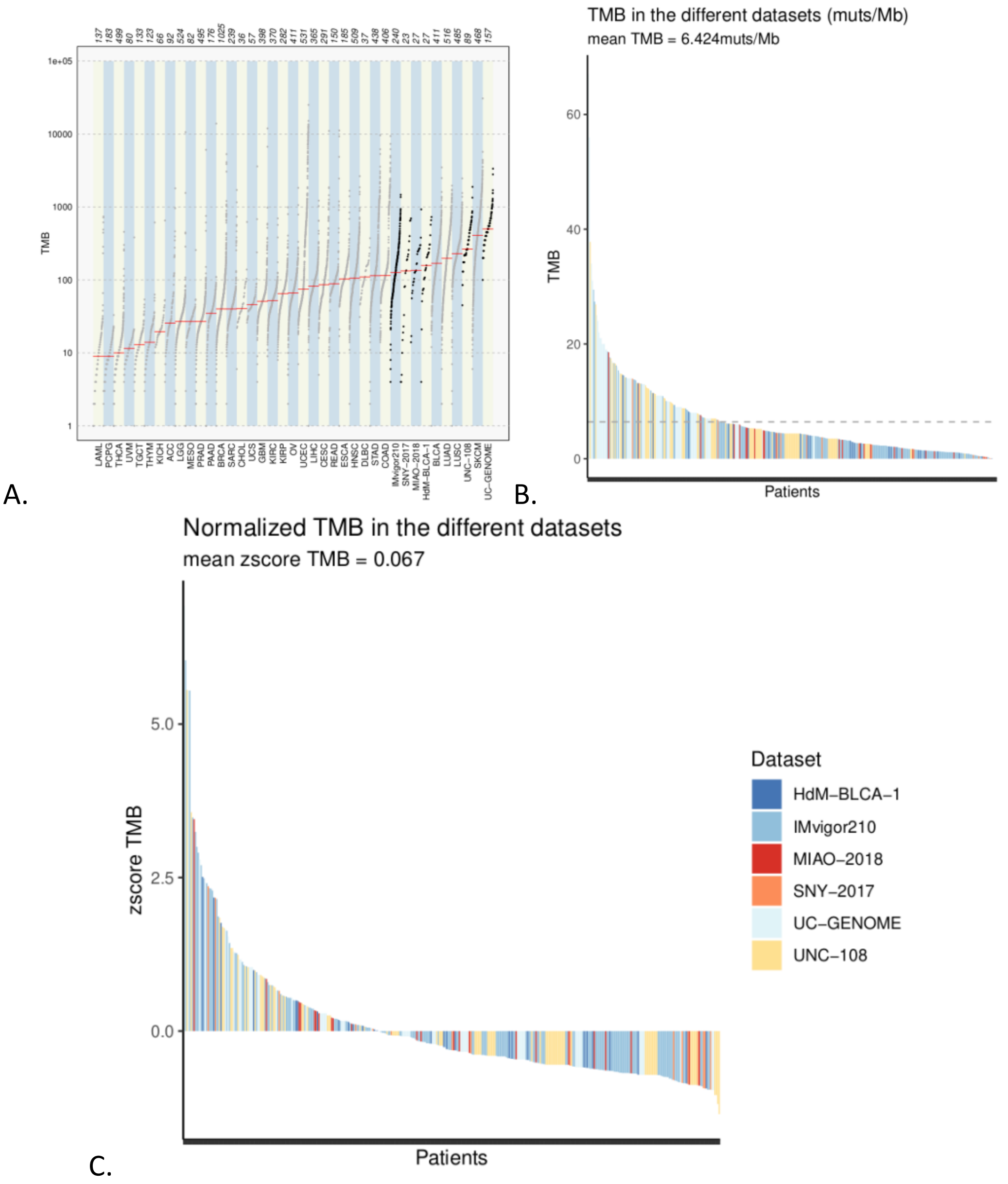


**Sup. Figure 16. Normalization of TMB values. A.** Distribution of TMB in the different datasets compared to other TCGA cohorts. The four datasets with WES data (IMvigor210, SNY-2017, MIAO-2018 and HdM-BLCA-1) show a similar number of mutations as the TCGA bladder cancer cohort (BLCA). The two datasets with panel DNA (UC-GENOME and UNC-108) show overall higher TMB estimates. **B.** TMB values sorted descendingly and coloured by dataset. **C.** Sorted z-score TMB coloured by dataset.

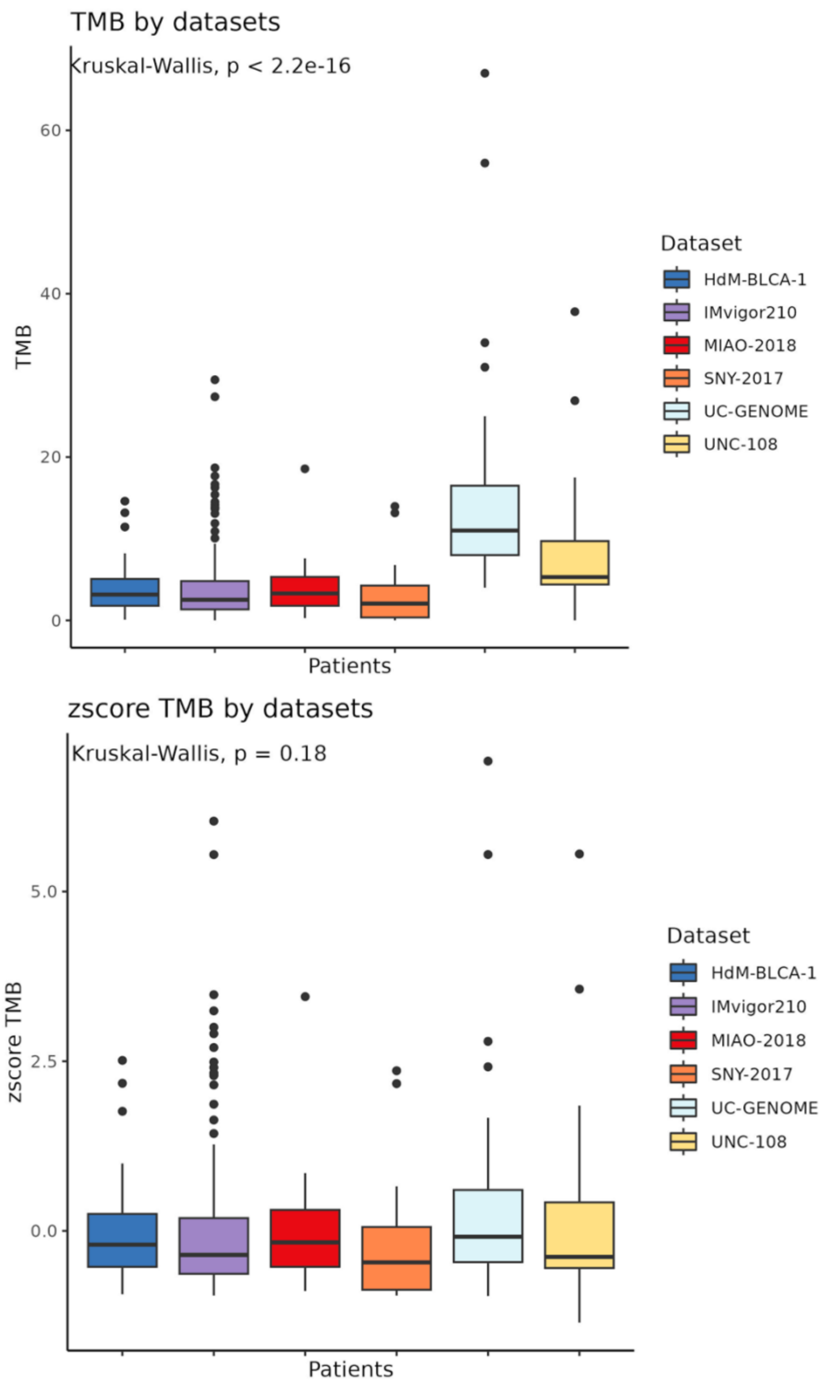


.

**Sup. Figure 17. TMB distribution before and after normalization. TMB by datasets shows** the distribution of TMB values in different datasets, there are significant difference among them. Zcore TMB by datasets shows the data after normalization; there are no differences among datasets.

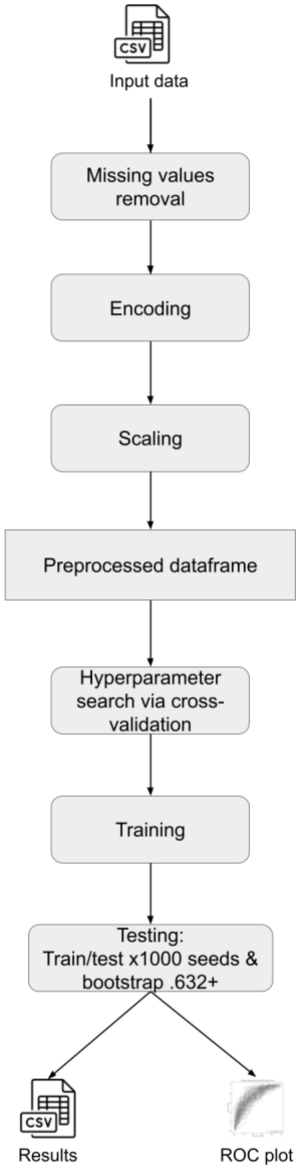


**Sup. Figure 18. Model flow chart.** The process of training and testing the model comprises multiple steps that start with data gathering followed by preprocessing steps and finally training and testing the model. A file containing all the results and a ROC plot are obtained.
